# METTL3 regulates exocytosis independently of m^6^A

**DOI:** 10.1101/2025.05.26.656168

**Authors:** Margalida Esteva-Socias, Devi Prasad Bhattarai, Cyrinne Achour, Poonam Baidya, Kanchan Kumari, Gaia Fontanari, Kerstin Seier, Hudson Pace, Sandhya Malla, Shaochun Zhu, Eva Lundin, Cathrine Broberg Vågbø, Marta Bally, Vinay Swaminathan, André Mateus, Ruth Rodriguez-Barrueco, Andreas Pich, Francesca Aguilo

## Abstract

RNA modification pathways are often mis-regulated in various cancers, with *N*^6^-methyladenosine (m^6^A) having a pivotal role in cancer progression and metastasis. Methyltransferase-like 3 (METTL3), a core component of the m^6^A methyltransferase complex, functions not only as an m^6^A writer but also promotes tumorigenesis through m^6^A-independent mechanisms. Here, we show that METTL3 is mislocalized to the cytoplasm in breast cancer tumors from patients, contributing to the oncogenic phenotype. Cytoplasmic METTL3 interacts with EXOC7, a key regulator of exocytosis, promoting its stabilization. Additionally, METTL3 regulates m^6^A-dependent alternative splicing of *EXOC7*. Silencing *METTL3* impairs vesicle trafficking and the breast cancer secretome – effects that do not rely on its enzymatic activity but instead involve METTL3-mediated stabilization of EXOC7 and potentially other exocyst components. Furthermore, METTL3 knockdown impairs invadopodia formation, collagen matrix invasion, and focal adhesion morphology in vitro, while inhibition of METTL3 catalytic activity does not. Our findings uncover non-catalytic roles of METTL3 in regulating exocytosis and the cancer secretome.

## Introduction

RNA modifications have a crucial role in regulating gene expression. Among these, *N*^6^-methyladenosine (m^6^A) is the most prevalent internal modification in messenger RNA (mRNA) and is central to regulating the mRNA life cycle (*1*). The m^6^A methyltransferase (writer) complex, comprising METTL3, METTL14, WTAP and additional cofactors (*2, 3*), is responsible for proper deposition of m^6^A during transcription (*4*). METTL3, however, is not always confined to this canonical complex. In several model systems, the zinc finger protein 217 (ZFP217) sequesters METTL3, preventing the formation of the METTL3-METTL14 heterodimer, and thereby restricting the methyltransferase activity (*5, 6*). m^6^A is reversible, and the demethylation is mediated by the erasers AlkB homolog 5 (ALKBH5) and fat mass and obesity-associated protein (FTO) (*7*). The m^6^A mark is recognized by a range of reader proteins that influence the stability, splicing, translation, and localization of mRNAs (*7*). Proper regulation of m^6^A is essential for normal development (*8, 9*), and dysregulation of m^6^A homeostasis is associated with human diseases, including cancer (*9, 10*). Recent evidence suggests that in cancer cells, METTL3 can also be aberrantly localized to the cytoplasm, where it facilitates translation independently of its catalytic activity (*11–15*). These findings suggest that METTL3 contributes to tumorigenesis through both its catalytic and non-catalytic functions, in a context-dependent manner that remains to be fully elucidated.

Breast cancer is the most frequently diagnosed female malignancy and a leading cause of cancer-related death among females (*16*). It is a highly heterogeneous disease that often leads to the development of distant metastasis (*17*). Several studies have shown that RNA modification pathways are dysregulated in breast cancer and play a significant role in disease progression (*18–20*). As such, these modifications are emerging as potential biomarkers for both the diagnosis and prognosis of breast cancer.

In cancer progression, including breast cancer metastasis, exocytosis has a central role in releasing cytokines, growth factors, proteases, and exosomes that degrade the extracellular matrix (ECM) (*21*). Among these proteases, matrix metalloproteinases (MMPs) are particularly important for ECM degradation and for facilitating cancer cell invasion. The exocyst complex, an evolutionarily conserved octameric protein complex composed of the EXOC1-EXOC8 subunits, is essential for this process, as it mediates exocytosis by tethering secretory vesicles to the plasma membrane (*22*). Within this complex, the EXOC7 subunit (also known as Exo70) has emerged as a key regulator of metastasis. EXOC7 interacts with the ARP2/3 complex, a central actin nucleator, to promote actin branching, lamellipodia formation, and directional cell migration (*23, 24*). It also contributes to invadopodia formation, specialized actin-rich protrusions that mediate ECM degradation during invasion (*25*). In breast cancer, the function of EXOC7 is particularly critical, as its knockdown markedly reduces cell invasiveness by impairing actin polymerization and blocking the secretion of MMPs (*25–28*). Moreover, *EXOC7* undergoes alternative splicing (AS), and isoform switching of *EXOC7* has been shown to regulate epithelial-mesenchymal transition (EMT) in breast cancer (*29*).

In this study, we reveal that METTL3 is mislocalized to the cytoplasm in breast tumors, where it interacts with EXOC7 and other subunits of the exocyst complex . We further show that *EXOC7* undergoes m^6^A-dependent alternative splicing. Silencing of METTL3 in breast cancer cells resulted in decreased vesicle trafficking and an impaired breast cancer secretome. Notably, these effects persist despite enzymatic inhibition, as they result from METTL3-mediated stabilization of EXOC7, revealing a previously unrecognized non-catalytic function of METTL3. Additionally, we show that METTL3 silencing, independent of its methyltransferase activity, impairs the formation of invadopodia and reduces the size of focal adhesion (FA)—structures that are necessary for extracellular matrix (ECM) remodeling, cell migration and invasion, further supporting the broader impact of METTL3 beyond m^6^A modulation.

Overall, these findings underscore the dual function of METTL3 in breast cancer, demonstrating that although its canonical role in m^6^A deposition remains critical, its non-catalytic function in regulating exocytosis is also important. By directly influencing vesicle trafficking, METTL3 not only fuels tumor growth and metastasis but also redefines our understanding of its impact beyond RNA modification. Given the fundamental role of vesicle trafficking in multiple cancer types, METTL3-mediated exocytosis likely represents a widespread oncogenic mechanism, suggesting its broader significance beyond breast cancer.

## Results

### METTL3 promotes breast cancer growth and metastasis

We have previously shown that silencing METTL3 leads to decreased proliferation, reduced colony formation, and increased apoptosis in breast cancer cell lines, indicating that METTL3 has a role in breast tumorigenesis (*30*). To further explore the oncogenic role of METTL3 in vivo, we injected METTL3-depleted and scramble control MDA-MB-231 cells into the mammary fat pads of athymic nude mice (fig. S1A). No significant differences in weight were observed between the two groups of mice over three weeks (fig. S1B). However, the xenograft tumors derived from METTL3 knockdown cells exhibited a significant reduction in weight and volume compared to controls (**Fig. 1, A** and **B** and fig. S1C). Consistent with these findings, immunohistochemistry (IHC) staining with Ki67 revealed a marked reduction in cell proliferation in the METTL3 knockdown tumors (**Fig. 1, C** and **D**).

**Fig. 1.**
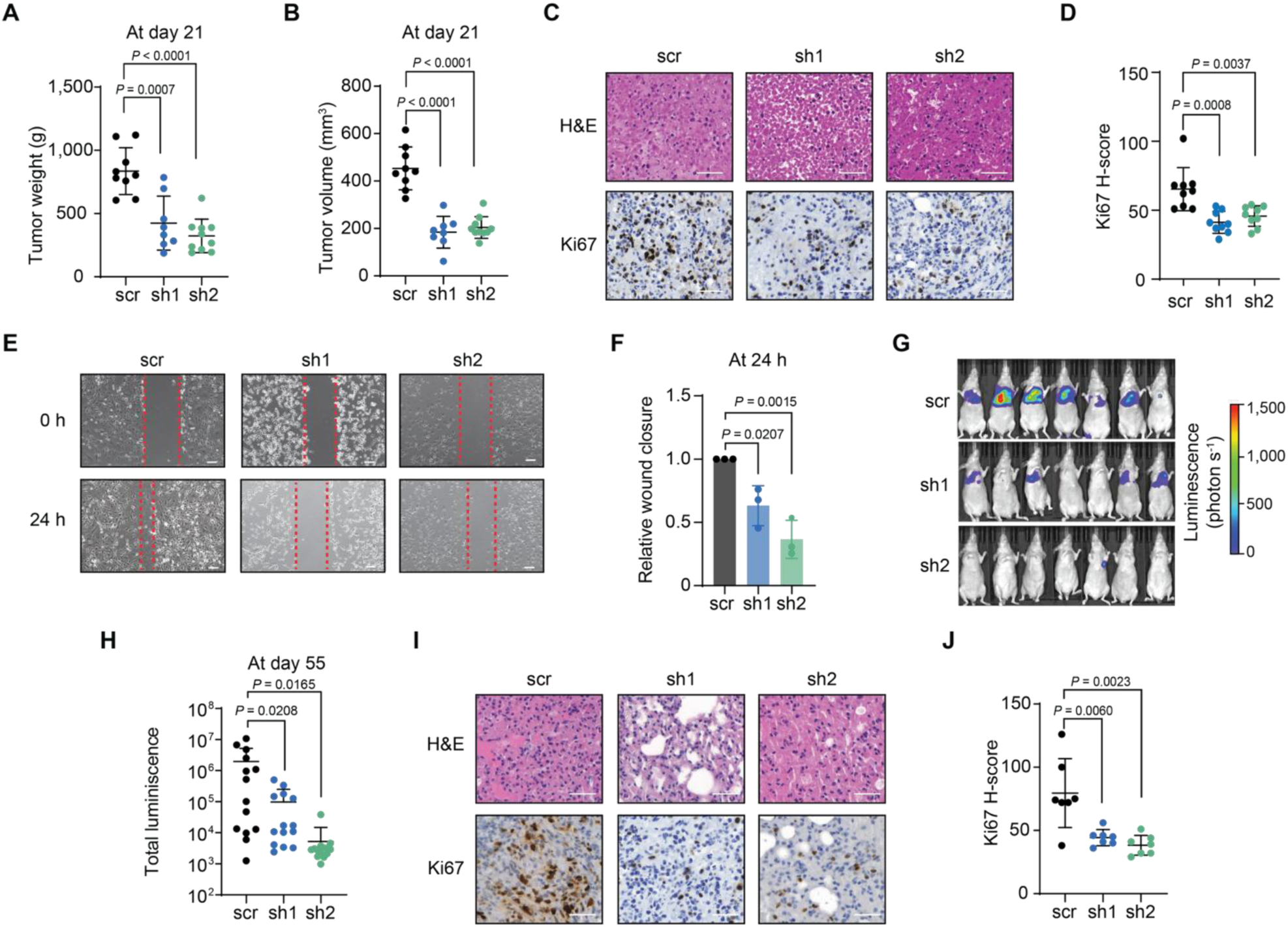
METTL3 promotes breast cancer growth and metastasis in vivo. (**A, B**) Tumor weight (**A**) and tumor volume (**B**) were measured in mice injected with scramble (scr) and METTL3 knockdown (sh1, sh2) MDA-MB-231 cells. (**C**) Haematoxylin and eosin (H&E, top) and Ki67 (bottom) staining of breast tissue sections from mice injected with scr or METTL3 knockdown MDA-MB-231 cells in the mammary fat pads. (**D**) Quantification of Ki67 staining in breast tissue sections from mice injected with scr or METTL3 knockdown MDA-MB-231 cells from (**C**). The intensity of the staining is associated with an H-score value (Methods). (**E**) Wound healing assay assessed in scr or METTL3-depleted MDA-MB-231 cells. (**F**) Bar graph illustrating relative wound closure at 24 h during the wound healing assay. (**G**) Bioluminescence imaging of mice injected with scr or METTL3 knockdown MDA-MB-231 cells. Scale represents the luminescence detected per pixel. (**H**) Quantification of total luminescence (sum for all the pixels) observed on the ventral and dorsal sides of the mice. (**I**) H&E (top) and Ki67 (bottom) staining of lung tissue sections from mice injected with scr or METTL3 knockdown MDA-MB-231 cells via the tail vein. (**J**) Quantification of Ki67 staining in lung sections from mice injected with scr or METTL3 knockdown MDA-MB-231 cells from (**I)**. The intensity of the staining is associated with a H-score value. Statistical analysis: Two-tailed Student’s *t*-test (**A, B, D, H,** and **J**) and one-way ANOVA with Dunnett’s correction for multiple comparison (**F**). Data are mean ± SD; *n = 9* (scr)*, 8* (sh1) and *10* (sh2) (**A**, **B**, and **D**), *n = 3* (**F**) and *n = 7* (**H** and **J**). Results are one representative of *n = 9* (**C**, **G** and **I**) and *n = 3* (**E**) independent biological experiments. Scale bars, 50 μm.

We next performed wound healing assays to investigate the metastatic potential of METTL3 in vitro. Silencing METTL3 led to a significant decrease in cell migration (**Fig. 1, E** and **F**). To assess its role in metastatic colonization in vivo, we performed experimental metastasis assays by injecting METTL3 knockdown and scramble control cells into the tail vein of athymic nude mice. METTL3 depletion delayed lung metastasis formation compared to controls (**Fig. 1, G** and **H**, and fig. S1, D and E). IHC staining with Ki67 in lung metastases further confirmed reduced proliferation in the METTL3 knockdown group (**Fig. 1, I** and **J**). In conclusion, these findings suggest that METTL3 promotes both breast cancer growth and metastasis in vitro and in vivo.

### METTL3 accumulates in breast cancer cell cytoplasm

Given the functional importance of METTL3 in promoting breast cancer growth and metastasis, we examined its expression in clinical samples. To this end, we performed IHC staining on a breast cancer tissue microarray (TMA) comprising 100 cases of invasive ductal carcinoma and 10 adjacent healthy epithelial tissues. Our analysis revealed that METTL3 was present in both the nucleus and the cytoplasm in breast cancer samples whereas it predominantly localized in the nucleus of healthy epithelial tissues (**Fig. 2A**). We further assessed staining intensity in these cellular compartments (fig. S2, A and B). Although the nuclear expression of METTL3 did not differ significantly between breast cancer and normal tissues, we observed an accumulation of METTL3 in the cytoplasm across all breast cancer subtypes that we analyzed (**Fig. 2, B** and **C**). Furthermore, we investigated the expression of METTL3 in a TMA containing normal healthy samples from various tissues. In these tissues, METTL3 predominantly localized to the nuclei, except in the bladder, lung and pancreas, where it also showed cytoplasmic accumulation (**Fig. 2D**).

**Fig. 2.**
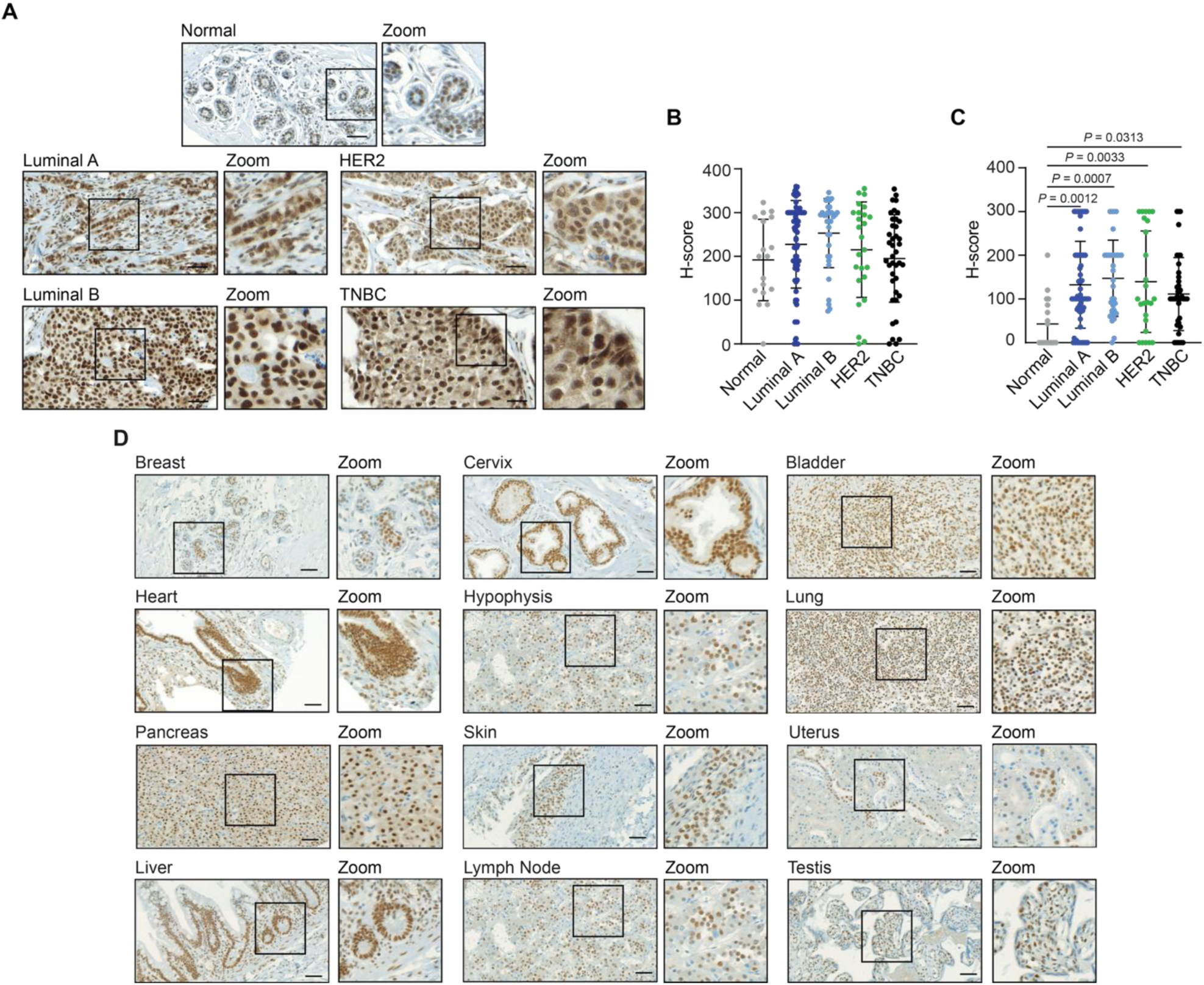
METTL3 is mislocalized in breast cancer cells. (**A**) IHC staining of METTL3 in normal epithelial tissue and breast cancer tissues from patients. (**B** and **C**) Quantification of IHC detection of METTL3 in the nucleus (**B**) and the cytoplasm (**C**) from (**A**). (**D**) IHC staining of METTL3 in various normal healthy tissues, including breast, cervix, bladder, heart, hypophysis, lung, pancreas, skin, uterus, liver, lymph node, and testis. Statistical analysis: One-way ANOVA with Dunnett’s correction for multiple comparison (**B** and **C**). Data are mean ± SD; *n > 18* (**B** and **C**). Results are representative of *n ≥ 18* (**A**) and *n = 6* (**D**). Scale bars, 50 µm.

To further investigate the localization of METTL3, we examined nuclear and cytoplasmic subcellular fractions from breast cancer cells (MCF7 and MDA-MB-231) and normal breast epithelial cells (MCF-10A and hTERT-HME1) by Western blot and immunofluorescence analysis. Unlike the patient specimens, METTL3 was found in both the nuclei and cytoplasm of both cancerous and non-cancerous cell lines (fig. S2, C and D). Together, these findings indicate that METTL3 localizes to the nuclei of healthy breast tissues, and that cell culture models do not accurately reflect the normal distribution observed in vivo.

### METTL3 interacts with the exocyst complex

To gain insights into the function of cytoplasmic METTL3, we isolated cytoplasmic fractions from MCF7 cells and performed immunoprecipitation of endogenous METTL3 followed by liquid chromatography-tandem mass spectrometry (LC-MS/MS) (fig. S3A). Our analysis identified METTL3 (bait) and several known components of the writer complex, including METTL14, WTAP, and VIRMA (**Fig. 3A**, fig. S3B, and table S1). Notably, we also detected components of the exocyst complex, a conserved octameric complex that mediates the tethering of secretory vesicles to the plasma membrane for exocytosis (*31, 32*). Gene Ontology (GO) analysis of the cytoplasmic METTL3 interactome further supported the enrichment of terms related to both the exocyst complex and the m^6^A methyltransferase complex, with relevant biological processes including mRNA methylation, protein transport, and vesicle docking (fig. S3C).

**Fig. 3.**
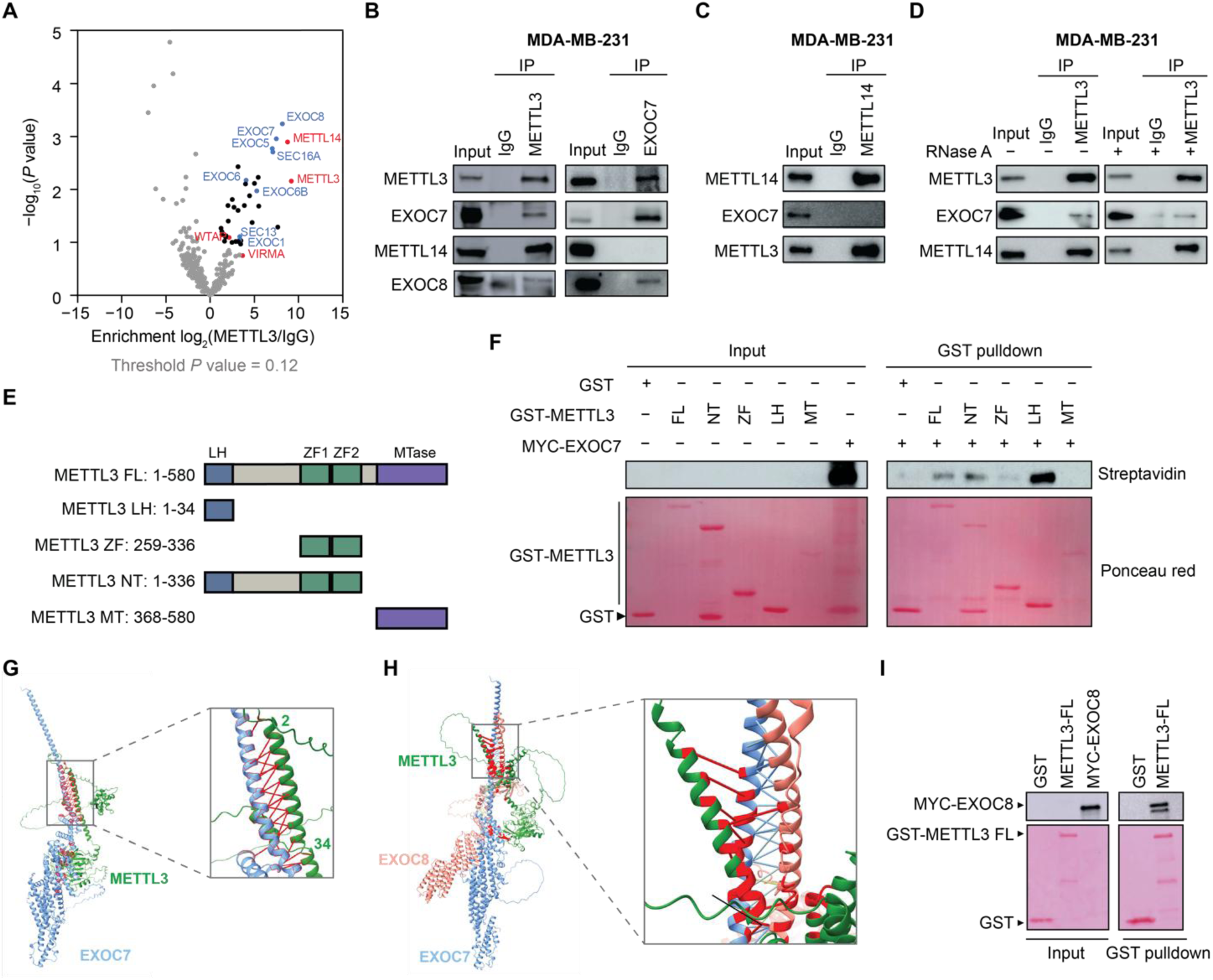
METTL3 interacts with subunits of the exocyst. (**A**) Volcano plot of METTL3-interacting proteins in the cytoplasm of MCF7 cells. *P* < 0.12 and log_2_ (METTL3/IgG) > 1 are used as thresholds. IgG is used as the negative control. The percentage of input used is 10%. (**B** and **C**) Immunoprecipitation (IP) of cytoplasmic extracts from MDA-MB-231 with antibodies against METTL3 and EXOC7 (**B**) and METTL14 (**C**) followed by immunoblotting with METTL3, EXOC7, EXOC8 and METTL14. IgG is used as the negative control. Input loaded is 10%. (**D**) Immunoprecipitation of cytoplasmic extracts from MDA-MB-231 pretreated with RNase A and with antibodies against METTL3 and EXOC7 followed by immunoblotting with METTL3, EXOC7 and METTL14. IgG is used as the negative control. Input loaded is 10%. (**E**) Schematic of GST–METTL3 purified proteins for in vitro pull-down analysis: full-length (FL), N-terminal (NT),zinc-finger (ZF),leader-helix (LH) methylase (MT). (**F**) GST pull-down assay with in vitro transcribed and translated MYC–EXOC7. Immunoblotting of MYC–EXOC7 was detected using Streptavidin–HRP. GST-tagged proteins were detected with Ponceau red staining. GST alone was used as the negative control. The percentage of input used for the MYC–EXOC proteins is 1% and 10% for the GST-tagged proteins. (**G**) Structural modeling of METTL3-EXOC7 interaction (left: overall model; right: Zoom onto LH domain interface. (**H**) Structural modeling of the ternary METTL3-EXOC7-EXOC8 interaction. (**I**) GST pull-down assay with in vitro transcribed and translated MYC–EXOC8. Immunoblotting of MYC–EXOC8 was detected using Streptavidin–HRP. GST-tagged proteins were detected with Ponceau red staining. GST alone was used as the negative control. The percentage of input used for the MYC–EXOC proteins is 1% and 10% for the GST-tagged proteins. Results are one representative of *n ≥ 3* (**B**, **F** and **I**) and *n = 2* (**C** and **D**) independent biological replicates.

Given its functional relevance in cell migration and invasion (*23, 26, 29, 30*), we focused on the interaction of METTL3 with EXOC7, one of the most enriched METTL3-interacting proteins identified. Using the MDA-MB-231 cell line, a well-established model for studying triple-negative breast cancer (TNBC), we found that both METTL3 and METTL14 localized in the nuclei and cytoplasm, while EXOC7 showed a similar distribution but with lower nuclear abundance (fig. S3D).

To confirm the interaction between METTL3 and EXOC7, we performed immunoprecipitation experiments of endogenous METTL3 in cytoplasmic fractions of MDA-MB-231 cells (**Fig. 3B**, left panel). Notably, in the reverse co-immunoprecipitation of endogenous EXOC7, we did not detect METTL14, suggesting that METTL3 may be part of distinct protein complexes, as previously suggested (**Fig. 3B**, right panel) (*33*). In line with this observation, we did not retrieve EXOC7 upon METTL14 immunoprecipitation (**Fig. 3C**). Furthermore, our analysis extended to a broader range of normal and cancer cell lines, where we previously assessed the cytoplasmic localization of METTL3 and EXOC7 (fig. S3, E to G). We observed interactions between METTL3 and EXOC7 in the ovarian cancer cell line OVCAR8, HEK293T, MCF-10A and hTERT-HME1 cells. However, no interaction was detected in mouse embryonic stem cells and U2OS cells, suggesting that the METTL3-EXOC7 interaction is context-dependent and may have varying biological roles depending on the cell type.

To further confirm the physical interaction between these proteins, we performed in vitro pull-down assays (fig. S3H). These experiments provided additional evidence supporting the direct interaction between METTL3 and EXOC7 and further revealed that the leader helix (LH) region of METTL3 is crucial for this interaction (**Fig. 3, E** and **F**). Although a weak EXOC7 signal was detected in the pull-down with the zinc finger domain construct, repeated assays using GST alone, full-length METTL3, and the zinc finger domain did not support a specific interaction with this region (fig. S3I). The LH of METTL3 has previously been shown to interact with WTAP (*34*), and AlphaFold modeling recapitulated this interaction (fig. S3J), albeit with modest confidence (ipTM = 0.22, pTM = 0.32).

We next employed AlphaFold modeling to gain structural insight into the METTL3–EXOC7 interaction. The model indicated that the METTL3 LH primarily engages the N-terminal α-helical region of EXOC7 (ipTM = 0.15, pTM = 0.44), with only a minor predicted contact at Asp324, consistent with experimental observations that the zinc finger region plays a limited role in binding. The predicted interface involves EXOC7 residues Val43–Gly107, forming a continuous α-helical surface complementary to the LH of METTL3 (**Fig. 3G**). Modeling of the ternary METTL3–EXOC7–EXOC8 complex further suggested that METTL3 also interacts with EXOC8 through its LH domain (**Fig. 3H**) (ipTM = 0.30, pTM = 0.38). This interaction was supported experimentally by immunoprecipitation of endogenous METTL3 followed by EXOC8 immunoblotting and GST pull-down assays (**Fig. 3, B** and **I**). In addition, co-immunoprecipitation of METTL3 also pulled down EXOC4 and EXOC5, further supporting interactions between METTL3 and multiple exocyst components (fig. S3K). Overall, these findings demonstrate that METTL3 interacts with the exocyst complex.

### m^6^A regulates *EXOC7* alternative splicing and EXOC7 protein stability

*EXOC7* isoform switching has been observed in specific cellular contexts (*29, 35, 36*). Based on exon 7 inclusion, *EXOC7* isoforms are classified into long isoforms (containing exon 7; isoforms 1, 4 and 6) and short isoforms (lacking exon 7; isoforms 2 and 5) (fig. S4A). *EXOC7* isoforms exhibit distinct expression patterns across various cell lines, with isoform 4 undetected in our cellular models (fig. S4B). The transition between short isoforms plays a role in EMT in breast cancer, with isoform 2 promoting EMT and metastasis, while isoform 5 reduces cell motility (*35*).

We previously demonstrated that METTL3 regulates *EXOC7* AS, reducing the abundance of the long isoform 6 and favoring the expression of the short isoform 5 upon METTL3 silencing, though we did not determine whether this regulation relies on its enzymatic activity (*30*). To address this, we first assessed whether *EXOC7* mRNA is m^6^A-modified. Analysis of publicly available MeRIP-seq datasets from MDA-MB-231 cells revealed a putative m^6^A site within exon 6 of *EXOC7* (fig. S4C) (*37*). Consistent with this observation, GLORI datasets in HEK293T and HeLa cells confirmed the presence of this m^6^A site immediately upstream of exon 7 (fig. S4, D and E) (*38*). Comprehensive mapping further revealed multiple m^6^A sites located within the 3′UTR of *EXOC7*, with one intronic site and two exonic sites, including exon 6 (fig. S4, D and E). Notably, the methylation level at these sites decreases following METTL3 inhibition or knockdown (fig. S4, D and E). In line with our prior findings, METTL3 depletion altered *EXOC7* splicing, leading to a marked reduction in the long isoform 6 and a modest increase in isoform 5 expression (**Fig. 4A**).

**Fig. 4.**
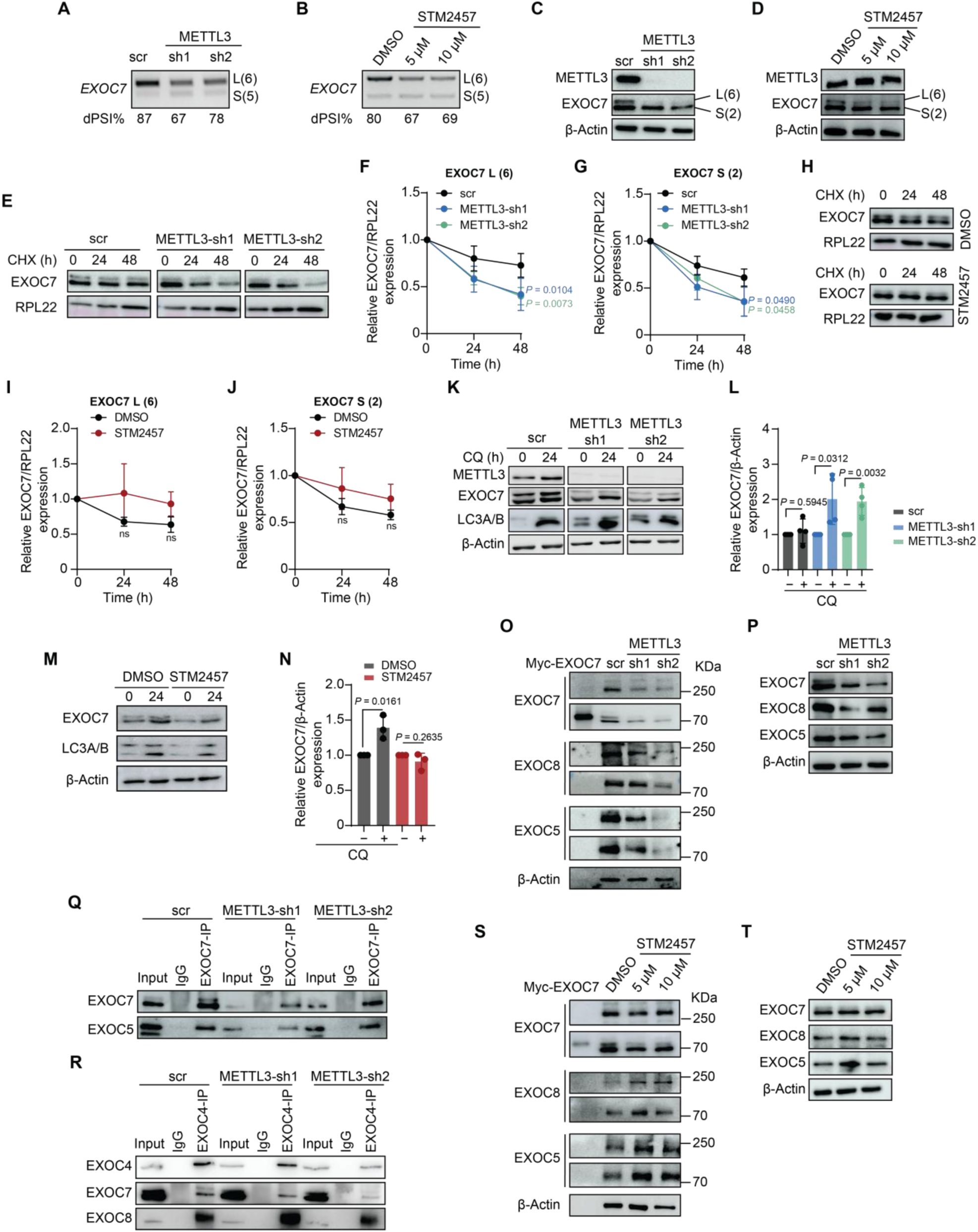
METTL3 regulates *EXOC7* alternative splicing and the exocyst function. (**A**) RT–PCR showing *EXOC7* exon-skipping in scramble (scr) and *METTL3* knockdown (sh1, sh2) cells. dPSI% indicates exon inclusion changes. (**B**) RT–PCR of *EXOC7* exon-skipping after STM2457 treatment. dPSI% shown. (**C**, **D**) Western blot of METTL3 and EXOC7 in scr and METTL3 knockdown MDA-MB-231 cells (**C**) and following STM2457 treatment (**D**). (**E**) Western blots of EXOC7 in scr and METTL3 knockdown HEK293T cells during CHX treatment. RPL22 as loading control. (**F** and **G)** Degradation curves of EXOC7 long (**F**) and short (**G**) from (**E**). *P* values at 24 h: long—sh1 = 0.0292, sh2 = 0.0363; short—sh1 = 0.0495, sh2 = 0.2470. (**H**) Western blots of EXOC7 from STM2457-treated HEK293T cells during a CHX treatment. RPL22 as loading control. (**I** and **J)** Degradation curves of EXOC7 long (**I**) and short (**J**) from (**H**). (**K**, **L**) Western blot and quantification of METTL3, EXOC7 and LC3A/B at 0 and 24 h after CQ treatment upon METTL3 depletion. β-Actin as loading control. (**M**, **N**) Western blot and quantification after STM2457 treatment. (**O** and **P**) Non-reducing PAGE (**O**) and Western blots (**P**) of EXOC7, EXOC8 and EXOC5 from Myc-EXOC7, scr and METTL3 knockdown MDA-MB-231 cells. β-Actin as loading control. (**Q** and **R**) Immunoprecipitation of cytoplasmic extracts using EXOC7 (**Q**) and EXOC4 (**R**) antibodies. IgG as control and 10% of input loaded. (**S** and **T**) Non-reducing PAGE (**S**) and Western blots (**T**) of EXOC7, EXOC8 and EXOC5 from Myc-EXOC7 and STM2457-treated cells. β-Actin as loading control. Statistical analysis: One-Way ANOVA with Dunnett’s correction for multiple comparison (**F** and **G**) and two-tailed Student’s *t*-test (**I**, **J** and **L**). Data are mean ± SD; *n ≥ 3* (**F**, **G**, **I**, **J**, **L** and **N**). All blots are one representative of *n = 3* independent biological replicates.

To further investigate whether this regulation depends on the m^6^A methyltransferase activity of METTL3, we conducted additional experiments using the specific METTL3 inhibitor, STM2457 (*39*). METTL3 inhibition in MDA-MB-231 cells was validated by measuring m^6^A levels on poly(A)-enriched RNA using mass spectrometry (fig. S4F). Additionally, pharmacological inhibition of METTL3 resulted in reduced cell proliferation and impaired wound healing (fig. S4, G and H). Notably, Western blot analysis and immunofluorescence staining revealed that STM2457 treatment slightly increased METTL3 protein levels and led to its predominant nuclear accumulation (fig. S4, I and J), potentially reflecting a compensatory mechanism in response to enzymatic inhibition. Similar to METTL3 silencing, treatment with METTL3 inhibitor also led to reduced expression of the long isoform 6 and increase of the short isoform 5, indicating that METTL3 controls *EXOC7* AS in an m^6^A-dependent manner (**Fig. 4B**).

To disregard that the reduced expression of the long isoform 6 resulted from altered RNA stability, we treated MDA-MB-231 cells with actinomycin D and measured *EXOC7* mRNA decay rates over time. No significant differences were observed following METTL3 knockdown or STM2457 treatment compared to controls (fig. S4, K and L), indicating that the decrease in long isoform 6 expression is not attributable to changes in mRNA stability.

We next focused on the short isoform 2 (instead of isoform 5) as it was the most abundant short isoform in MDA-MB-231 cells and detectable by Western blot (fig. S4M). For clarity, we will hereafter refer to isoform 6 as the long isoform and isoform 2 as the short isoform. When assessing EXOC7 protein levels, we observed a substantial decrease in the long isoform in METTL3 knockdown cells and inhibitor-treated cells, consistent with splicing defects, while the short isoform was slightly reduced only in the knockdowns (**Fig. 4, C** and **D**). To explore whether this reduction was linked to protein stability, we treated METTL3 knockdown cells with cycloheximide (CHX) and monitored EXOC7 protein levels over time. METTL3 depletion significantly shortened the half-life of both isoforms (**Fig. 4, E** to **G**), an effect not seen with enzymatic inhibition (**Fig. 4, H** to **J**).

We then examined whether METTL3 regulates EXOC7 protein stability by preventing its degradation through either the proteasomal or lysosomal pathways. Inhibition of the proteasome with MG-132 failed to restore EXOC7 protein levels in METTL3 knockdown cells (fig. S5, A and B). In contrast, lysosomal inhibition with chloroquine (CQ) in METTL3-depleted cells led to a marked accumulation of EXOC7, indicating that EXOC7 undergoes enhanced degradation through the autophagy-lysosome pathway in the absence of METTL3 (**Fig. 4, K** and **L**). However, STM2457 treatment did not recapitulate the effects of METTL3 silencing (**Fig. 4**, **M** and **N**), suggesting that this mechanism is independent of the methyltransferase activity of METTL3. This observation aligns with previous reports showing that EXOC7 undergoes autophagic degradation (*40*) and further supports a role for METTL3 in maintaining EXOC7 protein stability.

METTL3 has been shown to stabilize several of its binding partners, including METTL14, WTAP and p53, independently of its catalytic activity (*41–43*). Consistently, silencing METTL3 reduced the protein levels of other exocyst components that we found to interact with METTL3 (**Fig. 3B** and fig. S3K; fig. S5C to F), whereas treatment with the catalytic inhibitor STM2457 had no effect (fig S5, G to J).Given the role of METTL3 in EXOC7 protein stability, we next investigated exocyst complex assembly by performing non-reducing gel electrophoresis in scramble control and METTL3 knockdown cells. Immunoblotting for EXOC7, EXOC8 and EXOC5 revealed a decrease in both the oligomeric (migrating at ∼250 kDa) and monomeric (at ∼70 kDa) forms following METTL3 depletion (**Fig. 4O**). This decrease paralleled the reduction observed under reducing conditions (**Fig. 4P** and fig. S5, C to F), indicating that the diminished complex signal reflects reduced protein availability of the exocyst subunits rather than impaired assembly. Consistently, co-immunoprecipitation of multiple exocyst components demonstrated that the subunits remain associated and that complex formation is preserved in the absence of METTL3 (**Fig. 4, Q** and **R**). Moreover, treatment of MDA-MB-231 cells with STM2457 did not reduce either monomeric or oligomeric forms of the exocyst (**Fig. 4, S** and **T** and fig. S5, G to J). Altogether, these findings indicate that METTL3 regulates *EXOC7* AS through its catalytic activity and maintains EXOC7 protein stability via its scaffolding function, without disrupting the exocyst assembly; however, the overall amount of assembled complex to support vesicle trafficking might be compromised.

### METTL3 controls vesicle trafficking and secretion

Given that EXOC7 regulates the exocytosis of post-Golgi secretory vesicles at the plasma membrane (*44*), and that METTL3 depletion reduces the total abundance of assembled exocyst complexes, we sought to determine whether METTL3 influences vesicle trafficking and secretion. To this end, we monitored the transport of the transmembrane cargo protein vesicular stomatitis virus G protein (VSV-G) from the endoplasmic reticulum (ER) to the plasma membrane using the temperature-sensitive GFP-VSV-G^ts045^ construct (*45*). At 40°C, VSV-G is misfolded and retained in the ER; at 20 °C, it accumulates in the Golgi apparatus, and upon shifting to 32 °C, it is transported to the plasma membrane. To assess vesicle fusion with the plasma membrane, we performed immunostaining on non-permeabilized cells with the 8G5 monoclonal antibody, which specifically recognizes the extracellular domain of VSV-G. Quantifying and normalizing the surface VSV-G levels relative to total cellular levels revealed a 20-25% reduction in GFP-VSV-G^ts045^ incorporation into the plasma membrane following METTL3 silencing, suggesting that METTL3 is involved in VSV-G exocytosis (**Fig. 5A** and fig. S6A). Similar results were observed with EXOC7 knockdowns (fig. S6, B to E). However, treatment with the METTL3 inhibitor did not affect VSV-G secretion (**Fig. 5B** and fig. S6F).

**Fig. 5.**
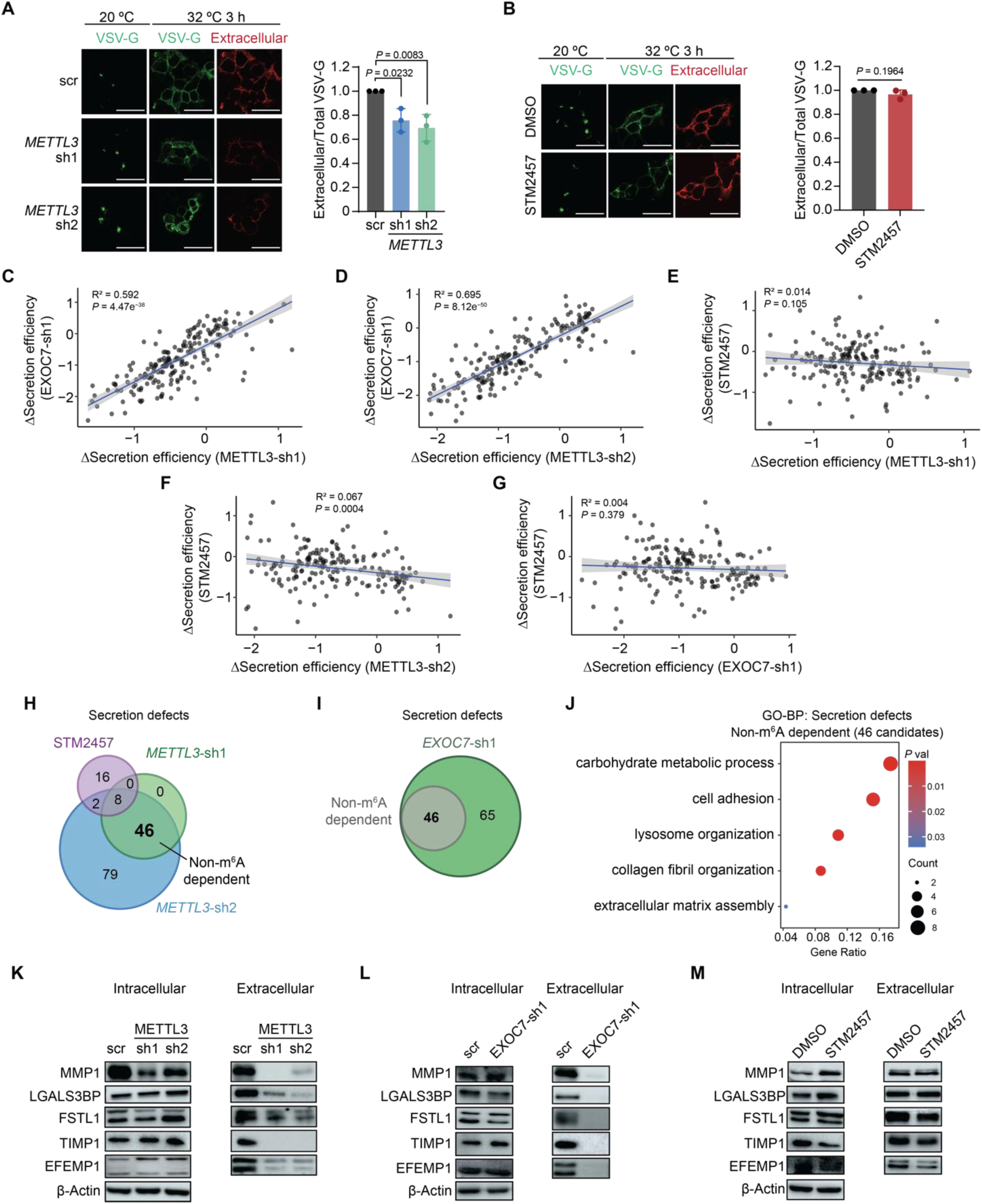
METTL3 is involved in vesicle trafficking regulation and modulates the secretome in MDA-MB-231 breast cancer cell line. (**A**) Left, immunofluorescence images of scramble (scr) and METTL3 knockdown (sh1 and sh2) HEK293T cells expressing GFP-VSV-Gts045 and collected after 3 h at 32 °C, then stained with the 8G5 antibody. Right, quantification shows the ratio of extracellular (8G5 staining) to total cellular VSV-G fluorescence. (**B**) Left, immunofluorescence images of HEK293T cells expressing GFP-VSV-Gts045 treated with DMSO or STM2457 after 3 h at 32 °C. Right, quantification shows the ratio of extracellular (8G5 staining) to total cellular VSV-G fluorescence. (**C** to **G**) Correlation of Δ Secretion efficiency between EXOC7-sh1 and METTL3-sh1 (**C**), EXOC7-sh1 and METTL3-sh2 (**D**), STM2457 and METTL3-sh1 (**E**), STM2457 and METTL3-sh2 (**F**) and STM2457 and EXOC7-sh1 (**G**). Proteins in common between different conditions were used to generate the plots. (**H**) Overlap between proteins that exhibit secretion defects in METTL3 knockdown compared to STM2457-treated MDA-MB-231 cells. (**I**) Overlap between downsecreted non-m^6^A-dependent proteins identified in (**H**) and EXOC7-sh1. (**J**) Gene ontology (GO) analysis of biological processes related to the common proteins showing a secretion defect identified in **(I**). (**K** to **M**) Intracellular and extracellular levels of secretory proteins in METTL3 knockdowns (**K**), EXOC7-sh1 (**L**) and STM2457-treated cells (**M**). Statistical analysis: One-way ANOVA with Dunnett’s correction for multiple comparison (**A**), two-tailed Student’s *t*-test (**B**) and Pearson correlation coefficients were calculated. R² values were obtained from simple linear regression (**C** to **G**). Data are mean ± SD; *n = 3* (**A** and **B**). Results are one representative of *n = 3* independent biological experiments (**A** and **B**, **K** to **M**). Scale bars, 30 µm.

Breast cancer cells secrete a diverse array of substances referred to as the secretome, which have been implicated in tumor development and invasion (*46–48*). Considering the negative impact of METTL3 silencing on vesicle trafficking, we investigated the secretome of scramble control and METTL3-depleted MDA-MB-231 cells by performing mass spectrometry analysis of the conditioned media, which was compared to the total proteome. We also conducted secretome analysis of EXOC7 knockdowns and cells treated with the METTL3 inhibitor STM2457. Protein hits were filtered to select secreted proteins, containing a signal peptide and without a transmembrane domain according to Uniprot database (*49*). Consistent with the defect in vesicle trafficking, we observed a reduction in protein secretion in both METTL3 and EXOC7 knockdowns compared to the scramble control (fig. S6, G to I and table S2). Notably, STM2457-treated cells showed only few proteins with decreased secretion (fig. S6J, table S3). This suggests that the methyltransferase activity of METTL3 has a limited role in regulating the MDA-MB-231 cell secretome. Moreover, while strong correlations were observed between the METTL3-dependent and EXOC7-dependent secreted proteins, this correlation was lost when comparing the secretomes of both METTL3 and EXOC7 knockdown cells to those of cells treated with the METTL3 inhibitor (**Fig. 5, C** to **G**).

To identify the m^6^A-independent secretory components, we next focused on proteins classified as showing secretion defects in both METTL3 and EXOC7 knockdowns but not in STM2457-treated cells. This analysis revealed 46 common proteins (**Fig. 5H** and fig. S6K). Strikingly, 100% of proteins with reduced secretion in the non-m^6^A-dependent secretome overlapped with those showing reduced secretion in EXOC7 knockdowns (**Fig. 5I** and fig. S6K). Gene ontology (GO) analysis revealed that these downsecreted proteins were enriched in biological processes crucial in cancer progression, including cell adhesion, collagen fibril organization, and ECM assembly (**Fig. 5J**). Several of these proteins, such as matrix metalloproteinase 1 (MMP1), LGALS3BP, FSTL1, TIMP1 and EFEMP1, have previously been identified in the breast cancer secretome regulating the metastatic ability of these tumors (*47, 50–53*). To further investigate these findings, we assessed the protein levels of the identified candidates in both cellular extracts and culture media. Consistent with the secretome analysis, METTL3 and EXOC7 knockdowns led to a significant reduction in their secretion (**Fig. 5, K** and **L**, fig. S6, L and M), while STM2457 treatment did not elicit the same effect (**Fig. 5M**). Although METTL3 depletion caused a modest reduction in intracellular MMP1 and EFEMP1, these proteins remained at appreciable levels, indicating that the secretion defect cannot be attributed to decreased protein synthesis.

Altogether, these findings suggest that METTL3 and EXOC7 jointly regulate the secretion of proteins involved in cancer progression, with METTL3’s function extending beyond its methyltransferase activity.

### METTL3 enhances ECM degradation for tumor invasion

Cancer cell invasiveness relies heavily on their ability to degrade the surrounding ECM. In METTL3 knockdown cells, proteins with reduced secretion – independent of m^6^A modification – were notably enriched in ECM organization categories. To explore this further, we performed a matrix degradation assay by plating scramble control and METTL3 knockdown MDA-MB-231 cells onto fluorescent-gelatin-coated coverslips. This assay evaluates invadopodia, F-actin-rich protrusions involved in ECM degradation and cancer cell invasiveness. Quantification of degradation areas showed that METTL3 knockdown cells had reduced invadopodia activity, reflecting impaired ECM degradation (**Fig. 6, A** and **B**). However, treatment with STM2457 did not have any effect on invadopodia activity (**Fig. 6, C** and **D**), suggesting that the m^6^A methyltransferase activity of METTL3 is not crucial for mediating ECM degradation.

**Fig. 6.**
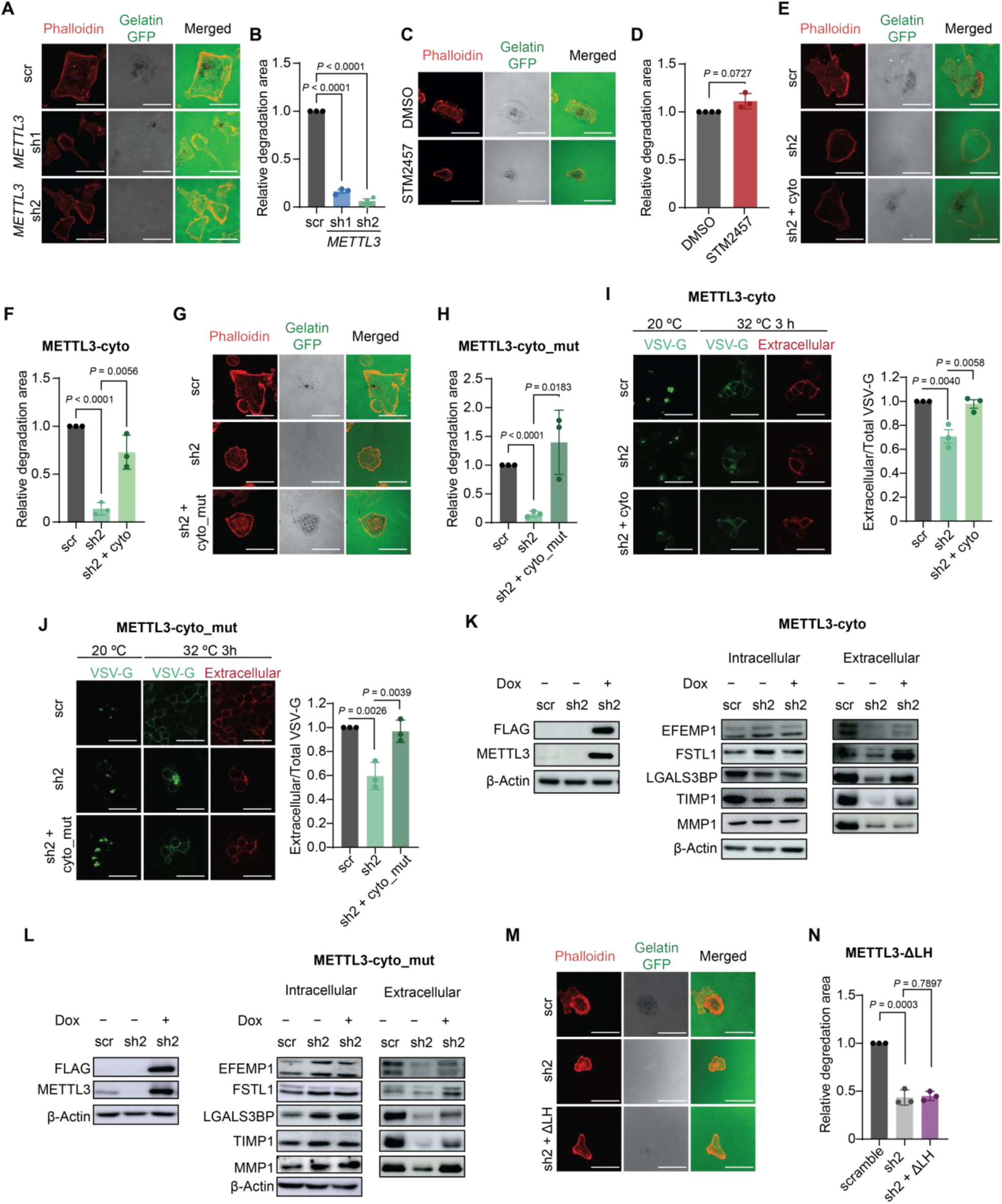
METTL3 promotes tumor cell invasion by facilitating ECM degradation through invadopodia formation. (**A**) Gelatin-degradation assay of MDA-MB-231 showing reduced invadopodia formation upon METTL3 depletion (sh1 and sh2) compared with control (scramble (scr)). Cells were cultured on Alexa-488 gelatin coated coverslips. Phalloidin is used as an F-actin marker. (**B**) Quantification of degradation areas in (**A**). (**C**) Gelatin-degradation assay of MDA-MB-231 upon STM2457 treatment. Cells were cultured on Alexa-488 gelatin coated coverslips. Phalloidin is used as an F-actin marker. (**D**) Quantification of areas of degradation in (**C**). (**E**) Rescue of invadopodia formation in METTL3 knockdown cells (sh2) by cytoplasmic METTL3 (sh2 + cyto). Phalloidin is used as an F-actin marker. (**F**) Quantification of areas of degradation in (**E**). (**G**) Rescue of invadopodia formation by catalytically dead cytoplasmic METTL3 (sh2 + cyto_mut). Phalloidin is used as an F-actin marker. (**H**) Quantification of degradation areas in (**G**). (**I** and **J**) Vesicle-trafficking rescue in HEK293T cells expressing GFP-VSV-G^ts045^ after 3 h at 32 °C, stained with 8G5 antibody; right panels show the ratio of extracellular to total VSV-G fluorescence for METTL3-cyto (**I**) and METTL3-cyto_mut (**J**). (**K** and **L**) Intracellular and extracellular levels of secretory proteins when rescuing METTL3 expression with METTL3-cyto (**K**) and METTL3-cyto_mut (**L**). (**M**) Invadopodia assay showing no rescue of invadopodia formation upon expression of METTL3–ΔLH (sh2 + ΔLH). Phalloidin is used as an F-actin marker. (**N**) Quantification of areas of degradation from (**M**). Statistical analysis: Two-tailed Student’s *t*-test (**B**, **D**, **F, H** to **J** and **N**). Data are mean ± SD; *n =*3 (**B**, **D**, **F, H** to **J** and **N**). Results are one representative of *n = 3* (**K** and **L**) independent biological replicates. Scale bars, 40 µm (**A**, **C**, **E**, **G** and **M**) and 30 µm (**I** and **J**).

To assess METTL3 specificity in regulating invadopodia formation, we engineered a doxycycline-inducible METTL3 cell line that cannot be targeted by shRNA2 and fused to a nuclear export signal (NES) (fig. S7, A and B). This enabled the depletion of endogenous METTL3 while maintaining cytoplasmic METTL3 expression (hereafter referred to as METTL3-cyto). Additionally, to dissect the catalytic from the non-catalytic functions, the catalytic center of METTL3 was mutated in the same cellular system (*aa395–398, DPPW → APPA*) (METTL3-cyto_mut) (fig. S7, A and B). As previously observed, METTL3 knockdown reduced EXOC7 protein levels, with a more pronounced effect on the long isoform (fig. S7, C to H). Reintroduction of both wild-type and mutant cytoplasmic METTL3 rescued the expression of the short EXOC7 isoform, consistent with its scaffold-mediated stabilization, while only wild-type METTL3 restored the long isoform, confirming the involvement of m^6^A for exon 7 inclusion of EXOC7 (fig. S7, C to H). METTL3-cyto_mut interacted with EXOC7, indicating that the intact catalytic domain of METTL3 is not required for such interaction (fig. S7I). Notably, treatment with the STM2457 inhibitor did not disrupt the interaction between EXOC7 and METTL3 (fig. S7J). Following endogenous METTL3 silencing, re-expression of either METTL3-cyto and METTL3-cyto_mut effectively rescued the invadopodia phenotypes (**Fig. 6, E** to **H**), confirming that the interaction between METTL3 and EXOC7, rather than its m^6^A methyltransferase activity, is necessary for this process. Moreover, both METTL3-cyto and METTL3-cyto_mut also restored vesicle trafficking defects in METTL3 knockdowns, suggesting that the cytoplasmic function of METTL3 contributes to exocytosis (**Fig. 6, I** and **J** and fig. S7, K and L). Reintroduction of METTL3-cyto and METTL3-cyto_mut also rescued the secretion of downsecreted proteins upon METTL3 silencing, including EFEMP1, FSTL1, LGALS3BP, and TIMP1, whereas MMP1 secretion was only recovered upon expression of the catalytically inactive METTL3 (**Fig. 6, K** and **L**).

To further investigate the role of METTL3-EXOC7 binding in regulating invadopodia formation, we examined the LH domain of METTL3, that we previously identified as critical for this association. We depleted endogenous METTL3 and introduced a METTL3 variant lacking the LH domain (METTL3-ΔLH), which was unable to interact with EXOC7 (fig. S7, M and N). Notably, re-expression of METTL3-ΔLH failed to restore invadopodia formation, highlighting the importance of METTL3-EXOC7 interaction in this process (**Fig. 6, M** and **N**).

Consistently, METTL3 silencing prevented the enhanced formation of invadopodia induced by overexpression of either the short or long EXOC7 isoform (fig. S8, A to F), an effect not observed following STM2457 treatment (fig. S8, G to J). Similarly, the formation of elongated tunneling nanotubes (TNTs), which mediate long-distance intercellular communication, was abolished when using an siRNA against METTL3 but persisted with STM2457 treatment (fig. S9, A to D). These findings confirm that METTL3 influences EXOC7 function through a catalytic-independent mechanism. Notably, overexpression of either the long or short EXOC7 isoform in the absence of METTL3 failed to induce invadopodia or elongated TNT formation, suggesting that METTL3 is an integral component of the complex required for these processes, although the underlying mechanism remains unclear. Alternatively, because METTL3 also affects the protein levels of other members of the complex, overexpression of EXOC7 alone may be insufficient to restore exocyst function.

We then generated tumor-derived spheroids - 3D cultures that mimic the in vivo cancer growth - and evaluated their ability to invade a collagen-rich ECM, simulating the tumor microenvironment. Notably, spheroid growth was completely abolished after 24 h of culture following METTL3 silencing (**Fig. 7, A** and **B**), but remained unaffected by METTL3 inhibitor treatment (**Fig. 7, C** and **D**), even after extended culture periods. Furthermore, METTL3 knockdown cells lost their ability to invade the collagen matrix (**Fig. 7, E** and **F**), while treatment with the STM2457 inhibitor resulted in only a modest, non-significant reduction in matrix invasion (**Fig. 7, G** and **H**).

**Fig. 7.**
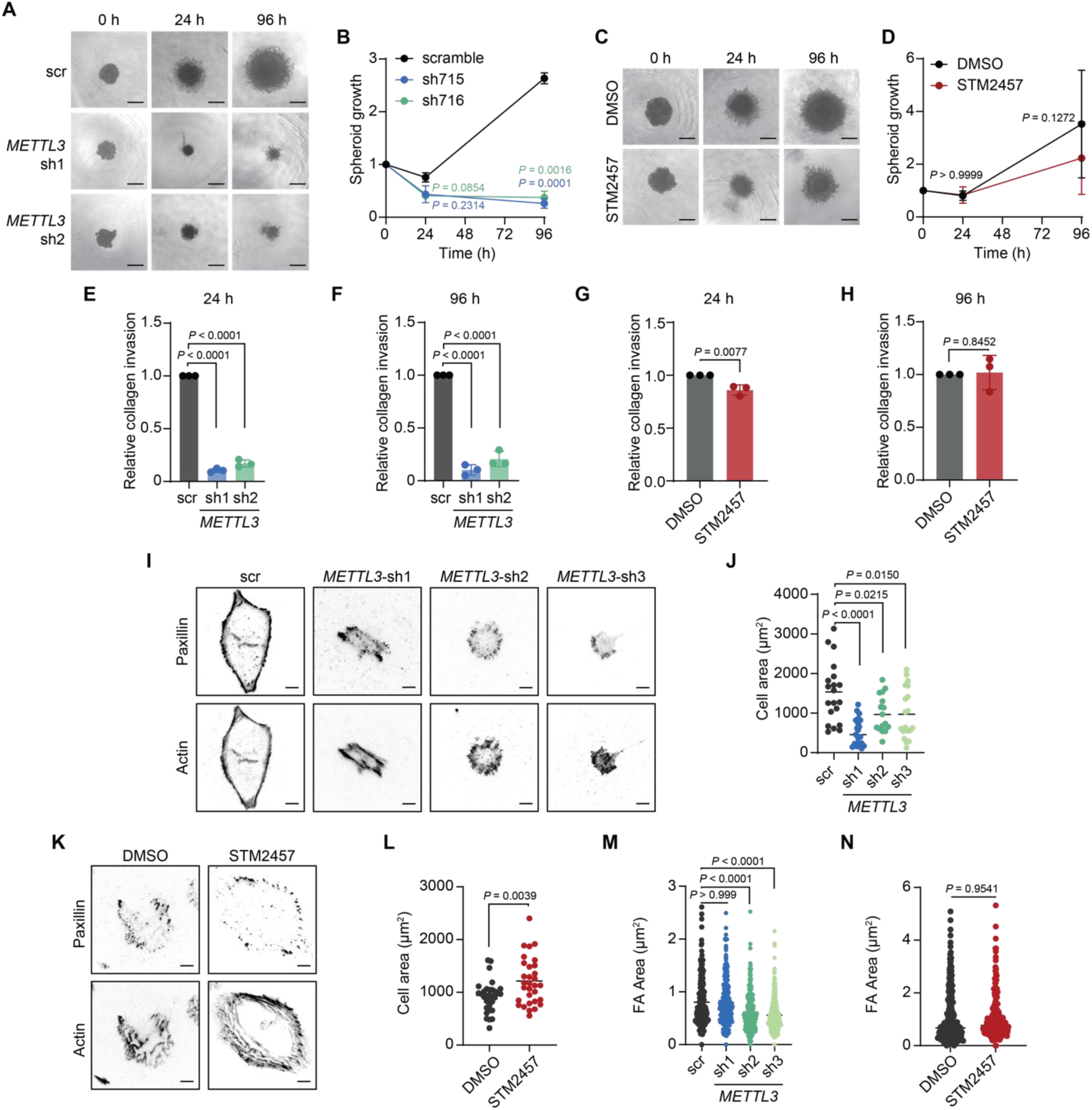
Functional impact of depletion and inhibition of METTL3 in invasion and cell adhesion. (**A**) Representative images of spheroids of scramble (scr) and METTL3 knockdown (sh1, sh2) MDA-MB-231 cells. (**B**) Spheroid growth measured after embedding scr or METTL3 knockdown MDA-MB-231 cells in Matrigel® at 24 and 96 h, relative to time 0 h (**C**) Representative images spheroids treated with DMSO or STM2457. (**D**) Spheroid growth measured after embedding DMSO or STM2457 treated cells in Matrigel® at 24 and 96 h, relative to time 0 h. (**E** and **F**) Quantification of invasive area at 24 h (**E**) and 96 h (**F**) of scr or METTL3 knockdown cells. (**G** and **H**) Quantification of invasive area at 24 h (**G**) and 96 h (**H**) of DMSO and STM2457-treated cells. (**I**) Representative images of scr and METTL3 knockdown on collagen I-coated glass, stained with Phalloidin-594 (Actin) and Alexa-488 (Paxillin). (**J**) Scatter plot quantification of cell area from (**I**). (**K**) Representative images of DMSO and STM2457-treated cells on collagen I-coated glass, stained as in (**I**). (**L**) Box plot quantification of cell area from (**K**). (**M** and **N**) Scatter plot of focal adhesion area of scr and METTL3 knockdown (sh1, sh2, sh3) cells (**M**) and STM2457-treated cells (**N**). Each point represents an average of 10 FAs per cell. Statistical analysis: Two-way ANOVA with Dunnett’s correction for multiple comparison (**B** and **D**), one-way ANOVA with Dunnett’s correction for multiple comparison (**E, F,** and **J**), two-tailed Student’s *t*-test (**G, H, L,** and **N**), Krustal-Wallis test (**M**). Data are mean ± SD; *n = 3* (**B**, **D**, **E** to **H**, **J**, **L**), *n ≥ 19* cells per condition (**M** and **N**). Results are one representative of *n = 3* (**A**, **C**, **I** and **K**) independent biological replicates. Scale bars, 200 µm (**A** and **C**) and 10 µm (**I** and **K**).

Cells sense and respond to ECM cues –such as composition, density, stiffness, viscoelasticity, and architecture– by generating forces at FA sites, which are transmitted downstream *via* mechanotransduction to regulate migration. To assess whether METTL3 depletion affects FA, scramble control, METTL3 knockdown, DMSO-treated, and STM2457-treated cells were seeded on collagen-coated coverslips and incubated for 24 h to allow FAs to form. Following incubation, cells were fixed and subjected to paxillin immunostaining, as paxillin is a key scaffold protein that localizes to FA, regulating their assembly, maturation, and signaling (*54*). Immunofluorescence analysis revealed substantial differences in cell morphology between scramble control and METTL3 knockdown cells, with the latter exhibiting a significantly reduced cell spread area (**Fig. 7, I** and **J**). In contrast, STM2457-treated cells displayed a modest increase in cell area compared to DMSO-treated controls (**Fig. 7, K** and **L**). However, when examining FA size, only shRNA2 showed a reduction relative to control cells (**Fig. 7M**). To address inconsistencies observed with the first two METTL3 shRNAs, a third shRNA targeting the 3’UTR of METTL3 was included, confirming a reduction in FA size. Notably, the pharmacological inhibition of METTL3 had no significant effect (**Fig. 7N**). Altogether these findings underscore the critical role of METTL3 in regulating the secretion of proteins associated with ECM invasion, ECM organization and cell adhesion, independent of its m^6^A methyltransferase activity.

## Discussion

In this study, we have uncovered a function of METTL3 in promoting exocytosis, extending its role beyond its well-established activity as an m^6^A methyltransferase. Abnormal expression of METTL3 and dysregulated m^6^A modification have been linked to various human diseases, including breast cancer (*18*). Here we provide evidence that METTL3 also contributes to breast tumorigenesis through m^6^A-independent mechanisms by regulating exocytosis via the exocyst complex.

Silencing METTL3 in breast cancer models resulted in impaired tumor growth and reduced lung metastasis, highlighting its critical role in breast cancer progression and metastasis. Additionally, we observed cytoplasmic accumulation of METTL3 in breast cancer tissuesfrom patients, whereas in normal breast epithelial tissues, METTL3 was predominantly nuclear, suggesting a shift in its subcellular localization during cellular transformation. Consistent with this, METTL3 is primarily localized in the nucleus in most normal tissues examined, except in the bladder, lung and pancreas-the latter two being secretory organs. Cytoplasmic localization of METTL3 has also been documented in other cancers, including lung cancer, chronic myeloid leukemia, and gastric cancer, where cytoplasmic METTL3 promotes translation independently of its catalytic activity (*12–14, 55*). However, its function in non-malignant cells remains poorly understood. Could the cytoplasmic redistribution of METTL3 in secretory organs be linked to exocytosis?

Aberrant protein localization is a well-established mechanism by which cancer cells promote transformation, survival, proliferation, and resistance to anti-neoplastic therapies (*56*). Recent studies have shown that acetylation of METTL3 facilitates its translocation to the cytoplasm and suppresses its chromatin binding (*57–59*). Beyond classical acetyltransferases such as p300, METTL3 acetylation is enhanced by the kinase PAK2, revealing a sophisticated crosstalk between phosphorylation and acetylation (*59*). Notably, PAK2 inhibition promotes ferroptosis, a form of programmed cell death driven by iron-dependent lipid peroxidation, and enhances cisplatin efficacy, in a manner dependent on METTL3 acetylation and its dissociation from chromatin (*59*). It would be of interest to determine whether the resulting effects on ferroptosis are not solely due to reduced m^6^A modification of enhancer RNAs and promoter-associated RNA, which stabilizes anti-ferroptotic transcripts, but also from impaired cytoplasmic localization of METTL3.

Our study uncovers a distinct cytoplasmic role for METTL3, expanding beyond its previously characterized function in translation. We identified METTL3 as an interacting partner of the exocyst complex, a key player in exocytosis which tethers secretory vesicles to the plasma membrane prior to fusion (*31, 32*). Given the established role of EXOC7 in breast tumorigenesis and its association with metastatic outcomes in TNBC (*29, 60*), we further characterized its interaction with METTL3. In addition to EXOC7, METTL3 also interacts with other exocyst subunits, suggesting that it may mediate exocyst function through interactions with additional members of the complex. Notably, METTL3–EXOC7 interaction occurs independently of METTL14, indicating that METTL3’s m^6^A methyltransferase activity is not required for binding EXOC7. Since METTL3 acetylation disrupts its interaction with METTL14 (*59*), it is plausible that the acetylated form of METTL3 constitutes the pool that interacts with EXOC7. Structurally, METTL3 binds EXOC7 through its N-terminal LH domain, which also interacts with the coiled-coil domain of WTAP. This interaction is known to mediate METTL3 nuclear localization for its canonical function (*2, 34*); therefore, binding to EXOC7 may retain a fraction of METTL3 in the cytoplasm to support exocyst-related processes.

EXOC7 undergoes isoform switching in breast cancer (*30*), a process that we demonstrate is regulated by m^6^A mRNA modification. Additionally, METTL3 stabilizes EXOC7 protein levels by protecting it from autophagy-mediated degradation, a mechanism previously reported to control EXOC7 turnover in ovarian cancer (*40*). Reduced EXOC7 levels upon METTL3 knockdown diminished exocyst complex assembly, as the complex could still form but was less abundant, leading to defective vesicle trafficking. Notably, this defect was not observed by catalytic inhibition, further supporting a non-enzymatic role of METTL3 in this process.

Our proteomics data align with these findings, revealing that METTL3 knockdown, but not catalytic inhibition, reduces the breast cancer secretome, particularly among proteins that are reliant on EXOC7. Previous studies have shown that METTL3 and METTL14 can transcriptionally regulate the senescence-associated secretory phenotype independently of m^6^A (*61*). Whether this holds true in breast cancer remains to be explored. Importantly, proteins diminished in the secretome upon METTL3 silencing are enriched in pathways critical for cancer progression, including cell adhesion, collagen fibril organization, and ECM assembly. Consistently, METTL3 silencing, without inhibiting its enzymatic activity, impaired invadopodia formation and affected collagen-rich ECM and FA architecture. Notably, even when EXOC7 was ectopically overexpressed in METTL3-depleted cells, defects in invadopodia and TNT formation persisted, indicating that METTL3 itself is essential for proper exocyst function and vesicle-mediated remodeling of the cytoskeleton. These observations suggest that METTL3 may also stabilize additional exocyst subunits with which it interacts, although this possibility warrants further investigation.

Current METTL3 inhibitors in clinical trials for advanced cancers primarily target its catalytic activity (*62*), which is dispensable for pro-tumorigenic functions like translation in lung and gastric cancers (*13, 14*). Our findings further demonstrate that the methyltransferase activity of METTL3 is not required for its function in exocytosis. Moreover, we observed an increase in METTL3 protein levels following inhibitor treatment, likely reflecting a compensatory response. Thus, approaches such as PROTAC-mediated degradation (*63*), which target METTL3 for proteolysis, may offer a more effective approach in contexts where cytoplasmic METTL3 drives oncogenic traits through non-catalytic mechanisms.

A key open question remains: Why is METTL3 involved in vesicle trafficking? While our findings indicate that the METTL3-EXOC7 interaction is RNA-independent, it is possible that these vesicles carry RNAs that, like proteins, are either secreted to the ECM or retained at the plasma membrane. One possibility is that METTL3 functions as a reader, binding to a specific subset of RNAs while simultaneously interacting with the exocyst complex, thereby ensuring the efficient export of these RNAs. The presence of extracellular mRNAs within extracellular vesicles (EVs) has been well-documented (*64–66*), yet the role of naked, non-vesicular extracellular RNA (exRNA) remains largely unexplored. Beyond intercellular communication, such non-vesicular exRNAs may contribute to RNA homeostasis or clearance. Furthermore, RNA-membrane interactions have been implicated in signal transduction, localized translation and cytoskeletal remodeling, processes that support cell surface plasticity and mesenchymal-like morphology in TNBC (*67–69*). Altogether, these observations support a broader view in which RNA turnover, translation, and vesicle trafficking maybe functionally interconnected, although further experimental evidence is required to substantiate this model.

Overall, our study provides novel insights into the role of METTL3 in exocytosis and its broader contribution to breast cancer. Targeting both the catalytic and non-catalytic functions of METTL3 could offer new avenues for diagnosis and treatment in breast cancer and other malignancies. Further investigations are warranted to unravel the complex interplay between the m^6^A-dependent and m^6^A-independent roles of METTL3 and to identify the mechanisms that govern its function in tumorigenesis.

## Materials and Methods

### Antibodies

The following antibodies were purchased from the indicated suppliers. For Western blot: anti-βACTIN (Sigma, A5441, 1:5,000), anti-METTL3 (Abcam, ab221795, 1:2,000), anti-METTL14 (Sigma, HPA038002, 1:4,000), anti-EXOC7 (Abcam, ab118792, 1:2,000), anti-EXOC5 (Santa Cruz Biotechnology, sc514802, 1:500), anti-EXOC8 (Abcam, ab254804, 1:2,000), anti-EXOC8 (Santa Cruz Biotechnology, sc-515532, 1:500), anti-EXOC4 (EMD Millipore, MABC570, 1:2,000), anti-FLAG (Sigma, F3165, 1:3,000), anti-GAPDH (Invitrogen, PA1-987, 1:2,000), anti-lamin A+C (Abcam, ab108992, 1:2,000), anti-HDAC1 (Abcam, ab19845, 1:2,000), anti-H3 (Abcam, ab8895, 1:8,000), anti-MMP1 (R&D Systems, MAB901-SP, 1:1,000), anti-RPL22 (Proteintech, 25002-1AP, 1:2,000), anti-TIMP1 (Proteintech, 16644-1-AP, 1:2,000), anti-LGALS3BP (Proteintech, 10281-1-AP, 1:2,000), anti-EFEMP1 (GeneTex, GTX111657, 1:2,000), anti-FSTL1 (Proteintech, 20182-1-AP, 1:3,000), anti-LC3A/B (Cell Signaling, 12741T, 1:2,000), Goat Anti-Mouse IgG H&L (HRP) (Abcam, ab6789, 1:10,000), Goat Anti-Rabbit IgG H&L (HRP) (Abcam, ab6721, 1:10,000). For co-immunoprecipitation experiments: anti-METTL3 (Bethyl, A301-567A, 3 µg), anti-EXOC7 (Abcam, ab118792, 3 µg), anti-FLAG (Sigma, F3165, 3µg), anti-METTL14 (Sigma, HPA038002, 3 µg), anti-EXOC4 (EMD Millipore, MABC570, 2.5 µg). For IF staining: anti-METTL3 (Bethyl, A301-567A, 1:100), anti-EXOC7 (Santa Cruz Biotechnology, sc-365825, 1:50), anti-FLAG (Sigma, F3165, 1:100), anti-VSVG (Kerafast, EB0010, 1:100), Phalloidin-iFluor 594 Reagent (Abcam, ab176757, 1:1,000), Phalloidin-iFluor 488 Reagent (Abcam, ab176753, 1:1,000) Alexa FluorTM 488 goat anti-mouse IgG (H+L) (Invitrogen, A11029, 1:1,000). Alexa FluorTM 568 goat anti-Mouse IgG (H+L) (Invitrogen, A110231, 1:1,000), Alexa FluorTM 568 goat anti-rabbit IgG (H+L) (Invitrogen, A11011, 1:1,000). For IHC stainings: anti-METTL3 (Abcam, ab221795, 1:200), anti-Ki67 (Abcam, ab16667, 1:100).

### Constructs and cloning strategy

To reconstitute METTL3 expression in MDA-MB-231, the cDNA of human *METTL3* was amplified with RevertAid First Strand cDNA Synthesis Kit and cloned into the doxycycline inducible pInducer20 lentivector (Addgene #44012), using Esp3I and XhoI as restriction enzymes. A nuclear export signal (NES) was added at the 3’end to enrich the expression of METTL3 to the cytoplasm. Silenced point mutations were generated at the shRNA2 targeting site to allow the inducible expression of METTL3. To produce catalytically dead METTL3, point mutations were generated at the catalytic site (DPPW ⟶ APPA). Additionally, to produce METTL3 lacking the LH domain, the N-terminal 1–34 amino acids were deleted.

To evaluate the interaction between METTL3 and EXOC7 *in vitro*, FLAG-tag and MYC-tag constructs were generated. Briefly, the cDNA of human METTL3 was amplified and cloned into pGEX6P-2 vector using BamHI and SalI as restriction enzymes, to further produce and purify GST-tagged METTL3 proteins in bacteria. The cDNA of human EXOC7 isoform 1 was amplified and cloned into pET22b vector using NdeI and EcoRI as restriction enzymes.

For EXOC7 overexpression and TNT formation assay, *EXOC7* short (isoform 2) and long (isoform 6) isoforms were cloned into pEGFP-C3-EXO70 vector (Addgene #53761). First, the isoform 1 was removed by digesting the vector using BamHI and XhoI as restriction enzymes. The cDNA of *EXOC7* isoforms 2 and 6 was amplified with RevertAid First Strand cDNA Synthesis Kit and cloned into the pEGFP-C3 empty vector using the restriction enzymes. To perform overexpression experiments followed by gelatin degradation assays, *EXOC7* isoforms were cloned into pCDNA3 vector. pEGFP-C3-*EXOC7*-isoform 2 and pEGFP-C3-EXOC7-isoform6 were used as a template to amplify the CDS of each isoform. Oligos used for the amplification contained a TCC sequence upstream followed by the restriction sites (BamHI for the forward primer and XhoI for the reverse). Both pCDNA3 and amplified isoforms were double digested with BamHI and XhoI, which were next ligated overnight at 16°C.

All constructs generated were confirmed by Sanger sequencing. All primers used for cloning purposes are described in table S4.

### Primers

All primers used in this study are listed in table S4.

### Cell culture

MCF7, MDA-MB-231, U2OS and HEK293T cell lines were cultured in Dulbecco’s Modified Eagle Medium (DMEM; Gibco) supplemented with 10% fetal bovine serum (Gibco) and 1% penicillin/streptomycin (Gibco). For MCF7 and MDA-MB-231 media was additionally supplemented with 10 µg.ml^−1^ insulin (Sigma-Aldrich). MCF10-A and hTERT-HME1 cell lines were cultured in DMEM/F12 (Sigma-Aldrich) supplemented with 5% heat-inactivated horse serum (Gibco), 20 ng.ml^−1^ epidermal growth factor (EGF; Sigma-Aldrich), 0.5 mg.ml^−1^ hydrocortisone (Sigma-Aldrich), 100 ng.ml^−1^ cholera toxin (Sigma-Aldrich), 10 µg.ml^−1^ insulin (Sigma-Aldrich), and 1% penicillin/streptomycin (Gibco). OVCAR-8 cell lines were cultured in RPMI 1640 medium (Gibco) supplemented with 10% fetal bovine serum (Gibco) and 1% penicillin/streptomycin (Gibco). CCE murine ESCs were culture and maintained on 0.1% gelatin-coated tissue culture plates under feeder-free culture conditions. The complete media composition consists of Dulbecco’s modified Eagle’s medium (DMEM; Gibco) 15% fetal bovine serum (Gibco), 1% MEM non-essential amino acids (Sigma-Aldrich), 0.1 mM of β-mercaptoethanol, 1% L-glutamine (Hyclone) and 1% penicillin/streptomycin (Gibco). Complete media was supplemented with 0.01 ng.µl^−1^ Leukemia inhibitory factor (LIF; R&D systems). All cell lines were cultured at 37°C under 5% CO_2_ in a humidified incubator. For METTL3 inhibition, cells were treated with STM2457 (MedChemExpress; HY-134836, 5 µM) for 48 h unless otherwise specified. To assess RNA stability, cells were treated with Actinomycin D (5 µg/ml) for specified times, followed by RNA extraction.

### Generation of knockdown cell lines

For lentivirus production, pLKO.1-shRNAs, the packaging vector pCMV-dR8.2 dvpr and the envelope vector pCMV-VSV-G (ratio 6:8:2) were co-transfected into HEK293T cells using Jet-PEI Polyplus following the manufacturer’s instructions. Culture media containing the lentiviral particles was collected after 48 and 72 h, filtered through a 0.45 µm filter and concentrated using Amicon Ultra-15 Centrifugal Filter (Merck). Cells were transduced with lentiviral supernatant and Polybrene (8 µg.ml^−1^). After 2 days, infected cells were selected with puromycin (1 µg.ml^−1^) for 4 days. The efficiency of METTL3 knockdown was evaluated by Western blot analysis. The shRNA sequences are listed in table S4.

### Generation and induction of METTL3-cyto, METTL3-cyto_mut and METTL3-ΔLH stable cell lines in MDA-MB-231

To reintroduce METTL3-cyto, METTL3-cyto_mut and METTL3-ΔLH, stable MDA-MB-231 cell lines were generated by lentiviral infection followed by selection with neomycin (8 mg.ml^−1^) for 8 days. After selection, exogenous METTL3 expression was induced with doxycycline (1 µg.ml^−1^) for 9 days. On day 9 of induction, non-induced cells were seeded for subsequent lentiviral infection with either a scramble control or METTL3 knockdown (sh2). Doxycycline-induced cells were infected with lentiviral particles targeting METTL3 (sh2 + dox, rescue). After 2 days, cells were selected with puromycin (1 µg.ml^−1^) for 4 days. The efficiency of METTL3 knockdown and rescue was confirmed by Western blot analysis. All downstream experiments were conducted after 4 days of puromycin selection.

### RNA extraction and RT–qPCR

Total RNA was extracted using the RNeasy Mini Kit (Qiagen) following the manufacturer’s instructions. 1 μg of total RNA was reverse transcribed using RevertAid First Strand cDNA Synthesis kit (Invitrogen) and quantitative PCR was performed using Power Up SYBR master mix (Applied Biosystems) on Quantstudio Flex6 instrument. *Β-Actin* was used for normalization.

### RT–PCR

1 μg of total RNA was reverse transcribed using the RevertAid First Strand cDNA Synthesis kit (Invitrogen). PCR was performed using the DreamTaq Green Master mix (Thermo Fisher Scientific) with specific primers to amplify the different splicing isoforms of *EXOC7*.

### mRNA purification

mRNAs were purified using Dynabeads following the manufacturer’s recommendations.

mRNAs were eluted twice with RNase-free water.

### mRNA mass spectrometry analysis

The RNAs were hydrolyzed to ribonucleosides by 20 U Benzonase® Nuclease (Santa Cruz Biotech, cat. no. sc-202391) and 0.2 U Nuclease P1 (Sigma, cat. no. N8630-1VL) in 10 mM ammonium acetate pH 6.0 and 1 mM magnesium chloride at 40 °C for 1 h. After that, ammonium bicarbonate to 50 mM, 0.002 U phosphodiesterase I and 0.1 U alkaline phosphatase (Sigma) were added, and incubated further at 37 °C for 1 h. The hydrolysates were mixed with 3 volumes of acetonitrile and centrifuged (16,000 x g, 30 min, 4 °C). The supernatants were dried and dissolved in 50 μl water for LC-MS/MS analysis of modified and unmodified ribonucleosides. Chromatographic separation was performed using an Agilent 1290 Infinity II UHPLC system with an ZORBAX RRHD Eclipse Plus C18 150 × 2.1 mm ID (1.8 μm) column protected with an ZORBAX RRHD Eclipse Plus C18 5 × 2.1 mm ID (1.8 μm) guard column (Agilent). The mobile phase consisted of water and methanol (both added 0.1% formic acid) run at 0.23 mL/min. For modifications, starting with 5% methanol for 0.5 min followed by a 2.5 min gradient of 5%–15% methanol, a 3 min gradient of 15%–95% methanol and 4 min re-equilibration with 5% methanol. A portion of each sample was diluted for the analysis of unmodified ribonucleosides which was chromatographed isocratically with 20% methanol. Mass spectrometric detection was performed using an Agilent 6495 Triple Quadrupole system, monitoring the mass transitions 268.1-136.1 (A) and 282.1-150.1 (m^6^A) in positive electrospray ionization mode.

### Whole-cell protein extraction

Cells were washed in cold PBS, pelleted, and lysed with ice-cold lysis buffer (50 mM HEPES pH 7.5, 150 mM NaCl, 3 mM MgCl_2_, 0.2% Triton X-100, 0.2% Nonidet NP-40, 10% glycerol) supplemented with protease inhibitor cocktail (Thermo Fisher Scientific) for 15 min on ice. Afterwards, lysates were sonicated (Bioruptor, Diagenode) for ten cycles with 30 s pulses on/off. Lysates were centrifuged at 13,300 rpm for 15 min, and supernatants were collected.

### Nuclear and cytoplasmic extracts

For nuclear and cytoplasmic extraction not involving co-immunoprecipitation experiments, a homemade extraction kit was used. Briefly, cells were washed with PBS and harvested by trypsinization or by scraping. Cells were pelleted by centrifugation at 2,200 rpm for 5 min at 4 °C, then resuspended in at least 5 volumes of cold cytoplasmic extraction buffer (10 mM HEPES pH 8, 10 mM KCl, 2 mM CaCl₂, 0.34 M sucrose, 10% glycerol, 0.2% IGEPAL CA-630, 1X Protease inhibitor cocktail). After vortexing, cells were incubated on ice for 10 min, followed by centrifugation at 8,000 rpm for 5 min at 4 °C. The supernatant was transferred to a new tube and kept as cytoplasmic extract. Nuclei pellets were then resuspended in 2.5 volumes of RIPA buffer (50 mM Tris pH 7.5, 100 mM NaCl, 3 mM EDTA, 0.5% IGEPAL CA-630, 1X Protease inhibitor cocktail). Samples were rotated for 15 min at 4 °C and sonicated for 10 cycles of 30 s on/off. Following centrifugation at maximum speed for 30 min at 4 °C, supernatant was collected and kept as the nuclear extract.

### Western blot detection of EXOC7 isoforms

EXOC7 protein isoforms were detected using a commercial anti-EXOC7 antibody (Abcam, ab118792), which recognizes an epitope common to all protein variants. The long isoform (isoform 6) and short isoform (isoform 2) were distinguished based on their migration patterns in SDS–PAGE, with the long isoform appearing at approximately 78–80 kDa and the short isoform at ∼74–75 kDa.

### CHX experiment

HEK293T cells were seeded at a density of 1.5 x 10^6^ cells in a 60 mm plate. After overnight incubation cells were treated with CHX at a final concentration of 100 µg.ml^-1^ for 0, 24 and 48 h, respectively. Cells were harvested for whole cell extracts on the indicated time points and proteins were subjected to immunoblotting. RPL22 was used as a loading control since β-actin has a shorter half-life than 48 h. Blots were quantified using Image Lab software.

### Proteasome and lysosome inhibition

MDA-MB-231 cells were seeded at a density of 3 x 10^5^ cells in 6 well-plate. After overnight incubation cells were treated with MG-132 (10 µM) for 10 h, or CQ (30 µM) for 24 h. Cells were harvested for whole cell extracts on the indicated time points and proteins were subjected to immunoblotting. Blots were quantified using Image Lab software.

### Non-reducing PAGE

The cultured medium was removed, and cells were resuspended in 1X PBS and centrifuged at 500 x g for 5 min. The cell pellet was then resuspended in NP-40 lysis buffer containing 10 mM Tris-HCl (pH 7.5), 150 mM NaCl, 0.5 mM EDTA, and 0.5% Nonidet™ P40 substitute, supplemented with 1X Halt protease inhibitor. The cell suspension was gently mixed by pipetting up and down and incubated on ice for 20 min before centrifugation at 17,000 x g for 15 min at 4 °C. The supernatant was collected as the whole cell extract.

Protein samples for non-reducing Western blot were prepared using 4X Laemmli buffer and separated on a 7.5% polyacrylamide gel without SDS. The running buffer contained 25 mM Tris, 192 mM glycine, and 0.1% SDS. Gels were run at 100V and transferred onto a nitrocellulose membrane at 80 V for 2 h under cold conditions. The membranes were blocked with 5% milk in PBS containing 0.1% Tween-20 (PBS-T) prior to immunoblotting with the indicated primary and corresponding secondary antibodies.

### Co-immunoprecipitation

Cytoplasmic extracts used for co-immunoprecipitation experiments were obtained using the NE-PER Nuclear and Cytoplasmic Extraction Reagents (Thermo Scientific). 500 µg of cytoplasmic extracts were pre-cleared with Pierce Protein G Agarose beads (Thermo Fisher) for 1 h at 4 °C and treated with rabbit IgG control or primary antibody (3 µg per immunoprecipitation) and incubated overnight at 4 °C. Next, washed Pierce Protein G Agarose beads (Thermo Fisher) were added, and the solution was incubated for an additional 3 h with gentle rotation. Then, beads were washed four times 5 min with ice-cold IP buffer (10 mM Tris pH 7.4, 1 mM EDTA, 1 mM EGTA pH 8, 150 mM NaCl, 1% Triton-X-100 and 0.2 mM Sodium Orthovanadate, 1X Protease inhibitor cocktail). Immunoprecipitated complexes were eluted using reducing sample buffer and analyzed by immunoblotting.

### Recombinant protein purification, *in vitro* transcription/translation and GST pull down

GST-tagged METTL3 proteins (full-length, N-terminal, methyltransferase, leader helix, or ZF domains) were purified from *Escherichia coli* strain Rosetta (DE3). Protein expression was induced overnight at 20 °C and cells were lysed by sonication and lysates were clarified by centrifugation and passed through a 0.45 µm filter. The lysates were incubated with Glutathione Sepharose 4B beads for 1.5 h at 4 °C. The resin was added to a gravity flow column and GST-tagged proteins were eluted with reduced L-Glutathione in elution buffer (125 mM Tris-HCl pH 8.0, 150 mM NaCl) after washing (125 mM Tris-HCl pH 8.0, 150 mM NaCl). GST-tagged proteins were stored in NETN buffer (20 mM Tris-HCl pH 8.0, 100 mM NaCl, 0.5 mM EDTA, 0.5% NP-40) and analyzed by SDS-PAGE followed by Coomassie blue staining.

In vitro transcription and translation for MYC-tagged EXOC7 was performed by incorporating biotinylated lysine to the translation reaction as a precharged ε-labeled biotinylated lysine-tRNA complex using Transcend Non-Radioactive Translation Detection systems (Promega). Briefly, 1 µg of pET22b -MYC-EXOC7 was incubated with TNT quick master mix and 1 µl methionine at 30 °C for 90 min.

To perform GST pull-down assay, MYC-EXOC7 protein and affinity purified GST-tagged METTL3 proteins were mixed in a ratio of 2.5:1 and incubated at 4 °C for 3 h. Then, equilibrated glutathione beads were added to this mixture and incubated at 4 °C for 2 h with gentle shaking. The protein complex was washed three times with NETN buffer (100 mM NaCl, 20 mM Tris-Cl pH 8.0, 0.5 mM EDTA and 0.5% (v/v) NP-40) and eluted with a loading buffer. Immunoprecipitated complexes were resolved by SDS-PAGE, transferred to PVDF membranes (Invitrogen), and detected with Streptavidin-HRP conjugate. For GST detection, the PVDF membranes were washed three times with PBST and incubated with 0.1% Ponceau red in acetic acid.

### Wound healing assay

2 x 10^5^ cells were seeded in a 12-well plate, and after 24 h cells were scratched to create a wound. After 24 and 48 h, pictures were taken, and the wound area was measured to assess cell migration. When STM2457 was used, the inhibitor was added when just after scratching at a final concentration of 5 µM.

### Cell proliferation

1 x 10^5^ cells were seeded in 6-well plate and were counted using trypan-blue (Bio-Rad) every second day for 6 days. When STM2457 was used, the inhibitor was added when seeding at a final concentration of 5 µM and maintained throughout the experiment.

### Immunofluorescence staining

Cells were fixed with 4% paraformaldehyde (Sigma-Aldrich) at room temperature (RT) for 15 min, washed three times, permeabilized for 30 min with 0.25% Triton-X-100 (Sigma-Aldrich) in PBS, and washed three times with PBS and blocked with 10% goat serum (Invitrogen) and bovine serum albumin (Hyclone) for 1 h. Cells were incubated with primary antibodies overnight at 4 °C and washed four times with PBS. Cells were incubated with secondary antibodies at RT. DAPI was used for nuclei staining. Imaging was acquired using Zeiss 710 and Leica SP8 X confocal microscope.

### Vesicle trafficking

Vesicle trafficking assays using GFP-VSV-G^ts045^ mutant were performed as described previously (*44, 70*). Briefly, cells were transfected with 1.5 µg of the VSV-G^ts045^-pEGFP-N1 vector using Lipofectamine™ LTX Reagent (Invitrogen). Cells were immediately placed at 40 °C, a restrictive temperature where the VSV-G mutant is misfolded and retained within the ER. Next, after overnight incubation, cells were shifted to 20 °C for 2 h for Golgi arrest and then moved at 32 °C and left for 1 and 3 h in the presence of 100 µg.ml^-1^ of cycloheximide (Sigma) to block protein synthesis. For each time point, cells were washed with PBS, fixed with 4% paraformaldehyde (PFA, Sigma-Aldrich) at RT for 15 min, washed and blocked for 15 min with blocking solution (1% bovine serum albumin, 10% goat serum). The coverslips were then incubated with the 8G5 monoclonal antibody against the extracellular domain of VSV-G (EB0010, Kerafast). No detergent was used in the immunofluorescence procedure. The amount of VSV-G fused to the plasma membrane was normalized to the total amount of VSV-G in the cells. For experiments performed using METTL3 inhibitor, cells were treated with 5 µM STM2457 for 48 h before seeding. STM2457 concentration was maintained throughout the experiment.

### Tunneling nanotube assay

8 x 10^4^ MDA-MB-231 cells were seeded in a 12-well plate. After overnight incubation, cells were transfected with 1 µg of plasmid (pEGFP, pEGFP-C3-EXOC7-short or pEGFP-C3-EXOC7-long) using Lipofectamine™ LTX reagent (Invitrogen). Images were captured 24 h after transfection using a Nikon ECLIPSE Ts2 inverted microscope coupled to a DMK NME33UX174 camera. TNT length was evaluated using ImageJ. When two cells were connected by a TNT, the total length of the projection was divided by two. For METTL3 silencing TNT experiments, MDA-MB-231 cells were treated with small interfering RNA (siRNA) targeting METTL3 or siRNA control for 72 h prior to TNT assay. For experiments with METTL3 inhibitor, cells were treated with 10 µM STM2457 for 48 h before seeding. STM2457 concentration was maintained throughout the experiment.

### Invadopodia formation assays

Invadopodia formation analyses were performed as previously described (*71*). Briefly, coverslips were coated with poly-L-lysine (50 µg.ml^−1^) for 20 min at 37 °C, crosslinked with 0.5% glutaraldehyde for 15 min at RT. After washing three times with PBS, coverslips were inverted on a 40 µl drop of 1 mg.ml^−1^green 488-labeled gelatin (Sigma-Aldrich) for 20 min at RT. Coverslips were then carefully washed with PBS and incubated with 5 mg.ml^−1^ of sodium borohydride to quench auto-fluorescence. Coverslips were incubated at least 1 h at 37 °C with complete medium before seeding. 1 x 10^5^ cells were seeded per coverslip and incubated 6 h at 37 °C. For METTL3 knockdowns, cells were seeded after 4 days of puromycin selection, while for METTL3 inhibitor after 48 h of treatment in the presence of 5 µM STM2457. Cells were fixed with 4% paraformaldehyde (Sigma-Aldrich) at RT for 15 min, washed three times with PBS, stained with Phalloidin-iFluor 594 (Invitrogen) and mounted on slides. Images were acquired using Leica SP8 X confocal microscope. Fluorescent gelatin degradation was quantified performed using ImageJ.

### Focal adhesions sample preparation

Coverslips were coated with 10 µg.ml⁻¹ PureCol® (Advanced Biomatrix) for 20 min at 37°C, washed with PBS, and blocked with 2% BSA in PBS for 1 h at 37 °C. After blocking, coverslips were washed twice with PBS, and 100,000 cells were seeded per condition and allowed to attach for 24 h. Following incubation, cells were washed twice with PBS and fixed with 4% PFA in cytoskeletal buffer (0.1 M MES at pH 6.1, 1.38 M KCl, 30 mM MgCl2, 20 mM EGTA) for 20 min at 37 °C. Cells were then incubated with Triton buffer (0.5% Triton in cytoskeletal buffer) at RT for 5 min, followed by incubation in glycine buffer (0.1 M Glycine in cytoskeletal buffer) for 10 min at RT with gentle rocking.

Subsequently, cells were washed three times with TBS (10 min per wash) and blocked with BTT buffer (4% w/v BSA, 0.1% v/v Tween-20 in 1X TBS) for 1 h at RT. Coverslips were incubated overnight at 4 °C with a primary antibody against Paxillin, diluted in BTT. The following day, cells were washed three times with TBS (10 min per wash) and incubated for 1 h at RT with a secondary antibody and phalloidin marker, diluted in BTT. Coverslips were then rinsed once with PBS and incubated for 5 min with 300 nM DAPI in PBS. Finally, coverslips were washed three times with TBS-T (0.1% v/v Tween-20 in 1X TBS) for 5 min each and stored in TBS at 4 °C until imaging.

### Total internal reflection microscopy (TIRFM)

The stained cells were imaged on an inverted Nikon Eclipse Ti-E2 microscope equipped with an Apo TIRF 60X oil-immersion objective (NA=1.49). TIRF images were acquired with laser excitation using the Ti-2 Lapp laser system with 488 nm and 561 nm wavelength. Epi fluorescence was acquired using a Spectra light source with 365 nm, 488 nm and 561 nm respectively. Images of the same cell were acquired in all 3 settings with 100ms exposure (TIRF 488, TIRF 561, Epi 365). For signal detection a Prime 95B sCMOS camera was used.

### Image analysis for focal adhesion quantification

Subsequent to image acquisition, the paxillin channel (TIRF 488) was used for quantification of focal adhesion morphology using the focal adhesion analysis server (FAAS) (*72*). A detection threshold of 4 and a minimum adhesion size of 4 pixels were used for all conditions. FA area and length for every 10 focal adhesions within a cell was then calculated and plotted as previously described (*73*).

### 3D invasion assay

The assay was conducted by initiating cell aggregation under different conditions. Briefly, 5 x 10^3^ cells (DMSO/ STM2457-treated and knockdown conditions) were seeded into low-attachment, U-bottom 96-well plates coated with 1% agarose. To initiate aggregation, the plates were centrifuged at 200 x g for 5 min, after which the cells were incubated for 5 days to form compact aggregates. On the same day, to assay collagenase activity, the aggregates in each well were embedded in a cold mixture of Matrigel ® Matrix (11573620, Fisher Scientific), 775 µg.ml^-1^ PureCol ® (Advanced Biomatrix), and DPBS that was mixed in 2:1:1 ratio. The plates were then briefly centrifuged at 200 x g and incubated at 37 °C and 5% CO_2_ to initiate Matrigel polymerization. Bright-field images were captured on day 0 and subsequently at 24 h and 96 h post-treatment. Thirty spheroids were analyzed per condition and biological replicate. For assessing spheroid growth over time, the total spheroid area was measured at day 0, while at days 1 and 4 (24 h and 96 h), only the core area, excluding the invasive region, was quantified. Spheroid invasion was calculated for each time point by subtracting the core area from the total area and normalizing to the initial spheroid area (day 0), using the following equation: *(Total area – Core area)/ Initial area (day 0)*. Image analysis was conducted using ImageJ software.

### Animal study

Female BALB/c nude mice (4 weeks old) were used in this study. For the tumor formation analysis, control and METTL3 knockdown MDA-MB-231 cells were injected into the mammary fat pad of mice. Mice weight was recorded every week until the day of sacrifice. Mice were sacrificed at day 21 and tumors were extracted. Tumor weight and volume were measured. For the metastasis xenograft model, MDA-MB-231 cells knockdown of *METTL3* expressing luciferase were injected in the mice tail vein. The metastatic spreading was monitored by bioluminescence on day 55. The mouse experiments were approved by the Ethical Committee for Animal Experimentation of the Catalan Government (project number #10568) were conducted in compliance with the guidelines for animal experimentation at IDIBELL, Universitat de Barcelona.

### Tissue staining and immunohistochemistry

Thin sections of formalin-fixed paraffin embedded tissues were deparaffinized, rehydrated and rinsed with water, and sections were mounted on slides. For H&E staining, hematoxylin and 0.5% eosin were used. For IHC, Ki67 and METTL3 antibodies were used. Staining intensity is recorded manually using the H-scoring system.

### TMA

Breast cancer tissue and multi normal human TMAs were purchased from US biomax (BC081116c and FDA999y) and stained as previously described in the above section. Staining intensity is recorded manually using the H-scoring system.

### H-score

Staining was quantified using an H-scoring system based on the percentage of tumor epithelial cells stained: [1 × (% cells 1+) + 2 × (% cells 2+) + 3 × (% cells 3+)]. The final score ranges from 0 to 300. Nuclear and cytoplasmic scorings were analyzed separately for the TMAs.

### Mass spectrometry analysis to identify METTL3 interacting proteins

500 μg of cytoplasmic extracts were pre-cleared with protein G magnetic beads (Bio-Rad) for 1 h at 4 °C then incubated overnight on a rotation with 3 μg of METTL3 antibody at 4°C. Protein G magnetic beads were added to the samples and incubated for 3 h at 4 °C with gentle rotation. Beads were washed four times with ice-cold immunoprecipitation buffer (10 mM Tris pH 7.4, 1 mM EDTA, 1 mM EGTA pH 8.0, 150 mM NaCl, 1% Triton and 0.2 mM Sodium Orthovanadate). Immunoprecipitated complexes were eluted with 5X reducing sample buffer. Following elution, immunoprecipitated complexes were mixed with 2 µl of 40% acrylamide solution for 20 min and resolved by SDS-PAGE. Gels were then stained with Coomassie blue (Thermo Fisher Scientific) and sent to the Research core unit proteomics, Medizinische Hohschule Hannover for mass-spectrometry. Briefly, gels were cut in slices, digested with trypsin, peptides were extracted and subjected to LC-MS analysis. Proteins were identified using MaxQuant Andromeda and human entries of Uniprot database (FDR < 0.01). Proteins quantified in at least three out of the four biological replicates were used. Missing values were imputed from a normal distribution with a down shift log_2_ of 1.8, and width 0.3. Significantly enriched proteins were identified using *t*-test; the enrichment threshold was Log2FC >1.

### Secretome sample processing and mass spectrometry analysis

After 4 days of puromycin selection for METTL3 and EXOC7 knockdown MDA-MB-231 cells or 48 h in the presence of 5 µM STM2457, two million MDA-MB-231 cells were seeded in a TC100 dish using complete DMEM media. The following day, the media was removed, and cells were washed two times with PBS (one quick wash and a second one for 2 min at RT) and incubated with 5 ml of FBS- and phenol red-free DMEM for 16 h. Afterwards, media was collected and filtered through a 0.45 µm filter and concentrated down to 300 µl using a 3 kDa Vivaspin™ 6 Columns (Cytiva) at 4,000 x g and 4 °C for 90 min. Samples were digested with a modified sp3 protocol, as previously described (*74, 75*). Briefly, samples were added to a bead suspension (10 μg of beads (Sera-Mag Speed Beads, 4515-2105-050250, 6515-2105-050250) in 10 μl 15% formic acid and 30 μl ethanol) and incubated shaking for 15 min at RT. Beads were then washed four times with 70% ethanol. Proteins were digested overnight by adding 40 μl of 5 mM chloroacetamide, 1.25 mM TCEP, and 200 ng trypsin in 100 mM HEPES pH 8.5. Peptides were eluted from the beads and dried under a vacuum. Peptides were then labelled with TMT10plex (knockdown experiment) or TMTpro (STM2457 experiment) (Thermo Fisher Scientific), pooled and desalted with solid-phase extraction using a Waters OASIS HLB Elution Plate (30 μm). Samples were fractionated onto 48 fractions on a reversed-phase C18 system running under high pH conditions, pooling every twelve fractions together. Samples were analyzed by LC-MS/MS using a data dependent acquisition strategy on a Thermo Fisher Scientific Vanquish Neo LC coupled with a Thermo Fisher Scientific Orbitrap Exploris 480. Raw files were processed with MSFragger (*76*) against a Homo sapiens FASTA database downloaded from UniProt (UP000005640) using standard settings for TMT. Protein hits were filtered using a custom R script to select proteins with a signal peptide without a transmembrane domain according to Uniprot database, as previously described (*49*). Data were normalized using vsn (*77*), and statistical significance was determined using limma (*78*) considering significant proteins as those with a log2 fold-change <-0.585 or log2 fold-change >0.585 and p-value < 0.05.

### Secretome data analysis

To quantify secretion-specific effects, we calculated a *secretion efficiency score* for each protein, defined as: *ΔSecretion efficiency = log₂FC(secreted) − log₂FC(full proteome).* This normalization allowed direct comparison of secretory output independently of overall expression changes. Based on this metric, proteins were classified into categories including *secretion defect*, *enhanced secretion*, *expression-driven/no secretion-specific change*, and *not significant*, according to the defined thresholds (table S5).

### Validation of secretome analysis

To validate the proteomic secretome analysis, conditioned media from each experimental condition were collected and concentrated to a final volume of 300 µl using 3-kDa molecular weight cut-off centrifugal filters. For Western blot analysis, 30 µl of each concentrated sample were mixed with reducing sample buffer, denatured, and loaded onto SDS–PAGE gel. The presence and relative abundance of selected secreted proteins were then assessed using specific antibodies, confirming the trends observed in the quantitative secretome data.

### Gene ontology (GO) analysis

Gene ontology (GO) analysis was performed using the web tool The Database for Annotation, Visualization and Integrated Discovery (DAVID) (https://davidbioinformatics.nih.gov/).

### Datasets

To study whether *EXOC7* is m^6^A modified, we used publicly available data for MeRIP-seq (GSE185494; https://www.ncbi.nlm.nih.gov/geo/query/acc.cgi?acc=GSE185494) and GLORI (GSE210563; https://www.ncbi.nlm.nih.gov/geo/query/acc.cgi?acc=GSE210563) from the GEO database.

### Statistical analysis

Data are shown as mean ± SD.GraphPad Prism 8.0.0 for Windows was used to perform the statistical analysis (GraphPad Software). The significance was determined using Student’s t-test, and two-Way ANOVA with Dunnett’s correction for multiple comparison

## Acknowledgments

The authors thank members of the Aguilo laboratory for useful discussion; the Biochemical Imaging Centre Umeå at Umeå University and the National Microscopy Infrastructure, NMI (VR-RFI 2023-00163) for providing assistance in microscopy; M. Lindberg for assisting with the cloning; S. Gildlund for performing IHC staining; and N. Nakamura for his generous gift of GFP-VSV-Gts045 plasmid.

## Author contributions

Conceptualization: FA.

Methodology: MES, DPB, CA, PB, KK, GF, KS, HP, SM, VS, AM, RRB, AP, FA.

Investigation: MES, DPB, CA, PB, KK, GF, KS, HP, SM, SZ, EL, MB, CBV, VS, AM, RRB.

Data curation: MES, CA, VS, AM, AP, FA.

Formal analysis: MES, DPB, CA, VS, AM, RRB, AP, FA.

Writing – original draft: MES, CA, FA.

Writing – review and editing: all authors.

Visualization: FA, MES, CA.

Supervision: FA.

Project administration: FA.

Funding acquisition: FA.

## Data and materials availability

The mass spectrometry proteomics data have been deposited to the ProteomeXchange Consortium via the PRIDE partner repository with the dataset identifiers, PXD066045 (https://www.ebi.ac.uk/pride/archive/projects/PXD066045) and PXD062959 (https://www.ebi.ac.uk/pride/archive/projects/PXD062959).

