## Supplementary Figures for "METTL3 regulates exocytosis independently of m^6^A"

Esteva-Socias M. *et al*

**Figure S1**

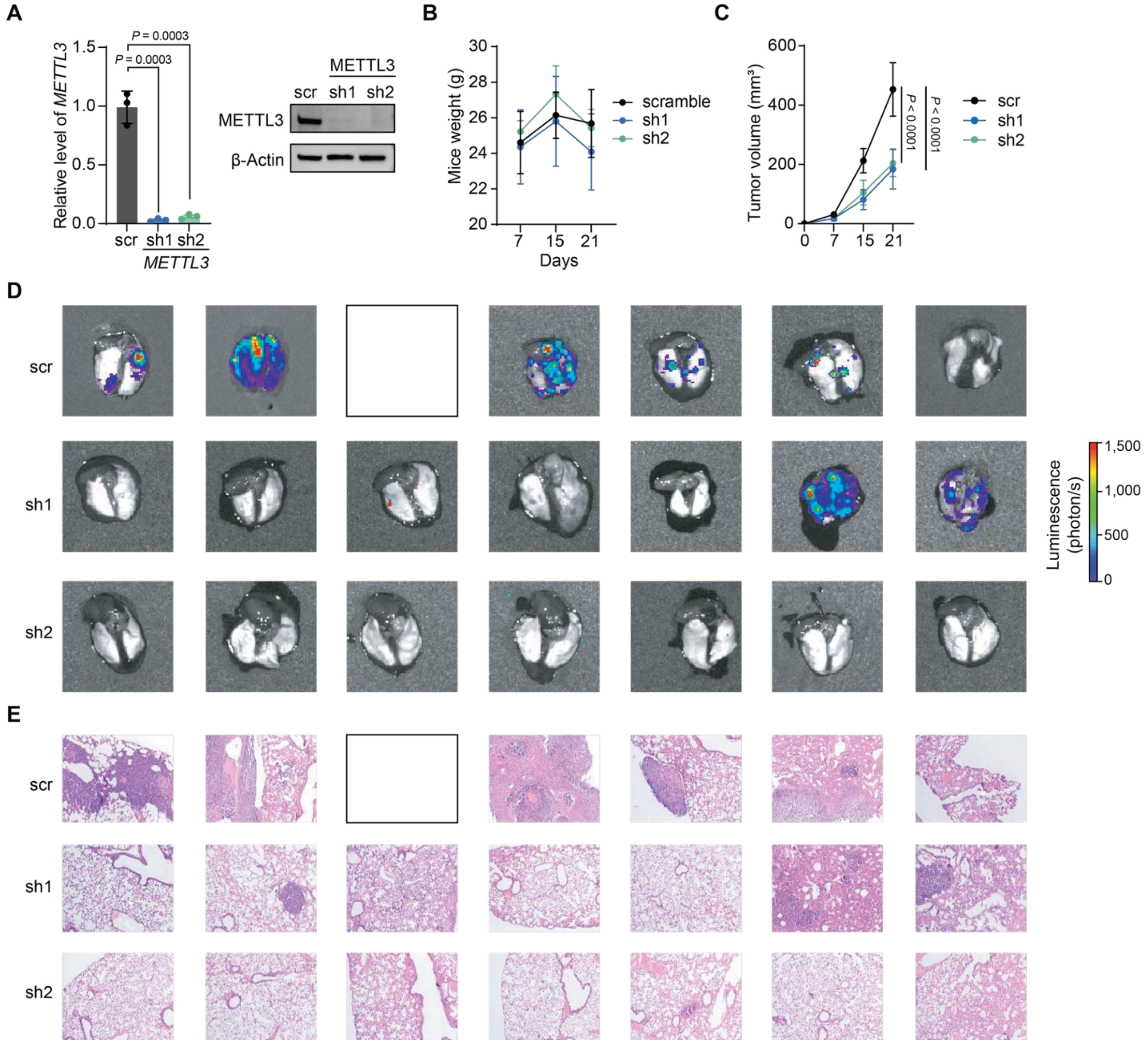


**Fig. S1. METTL3 promotes lung metastasis in mice.** (**A**) *METTL3* mRNA levels (left) and protein expression (right) upon depletion of METTL3 (sh1 and sh2) in MDA-MB-231 cells. (**B**) Mice weight and (**C**) tumor volume was measured over three weeks in mice injected with control (scramble (scr)) or METTL3 knockdown MDA-MB-231 cells (sh1 and sh2) in the mammary fat pads. (**D**) Bioluminescence image of lungs from mice injected with scr and METTL3 knockdown (sh1 and sh2) MDA-MB-231 cells *via* the tail vein. (**E**) Hematoxilin and Eosin (H&E) staining of lung sections from mice injected with scr and METTL3 knockdown (sh1 and sh2) MDA-MB-231 cells from (**D**). Statistical analysis: Two-tailed Student's *t*-test (**A** to **C**). Blank squares in (**D**) and (**E**) indicate data not available due to mice death before the study was completed. Data are mean ± SD; $n = 3$ (**A**) and $n = 9$ (**B** and **C**).

#### Figure S2

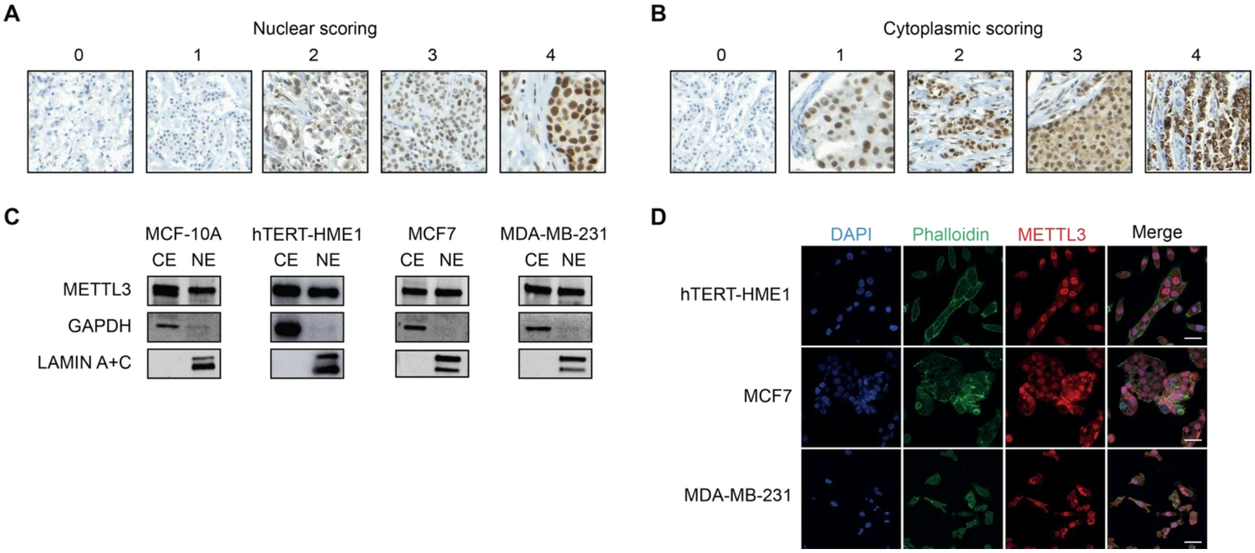


**Fig. S2. METTL3 localizes in both nuclei and cytoplasm of cell lines.** (**A, B**) IHC staining in normal epithelial and breast cancer tissues with low to high (0 to 4) expression of METTL3 in the nucleus (**A**) and in the cytoplasm (**B**) used as a reference for the H-score analysis. (**C**) Cellular fractionation of METTL3 in MCF-10A, hTERT-HME1, MCF7 and MDA-MB-231 cell lines. GAPDH and lamin A+C were used as loading controls for nuclear (NE) and cytoplasm (CE) extracts, respectively. (**D**) Immunofluorescence images of METTL3 in hTERT-HME1, MCF7 and MDA-MB-231 cell lines. DAPI and Phalloidin-488 were used as nuclear and cytoplasmic markers, respectively. Results are one representative of $n = 2$ independent biological experiments (**C** and **D**). Scale bars, 30 µm.

### Figure S3

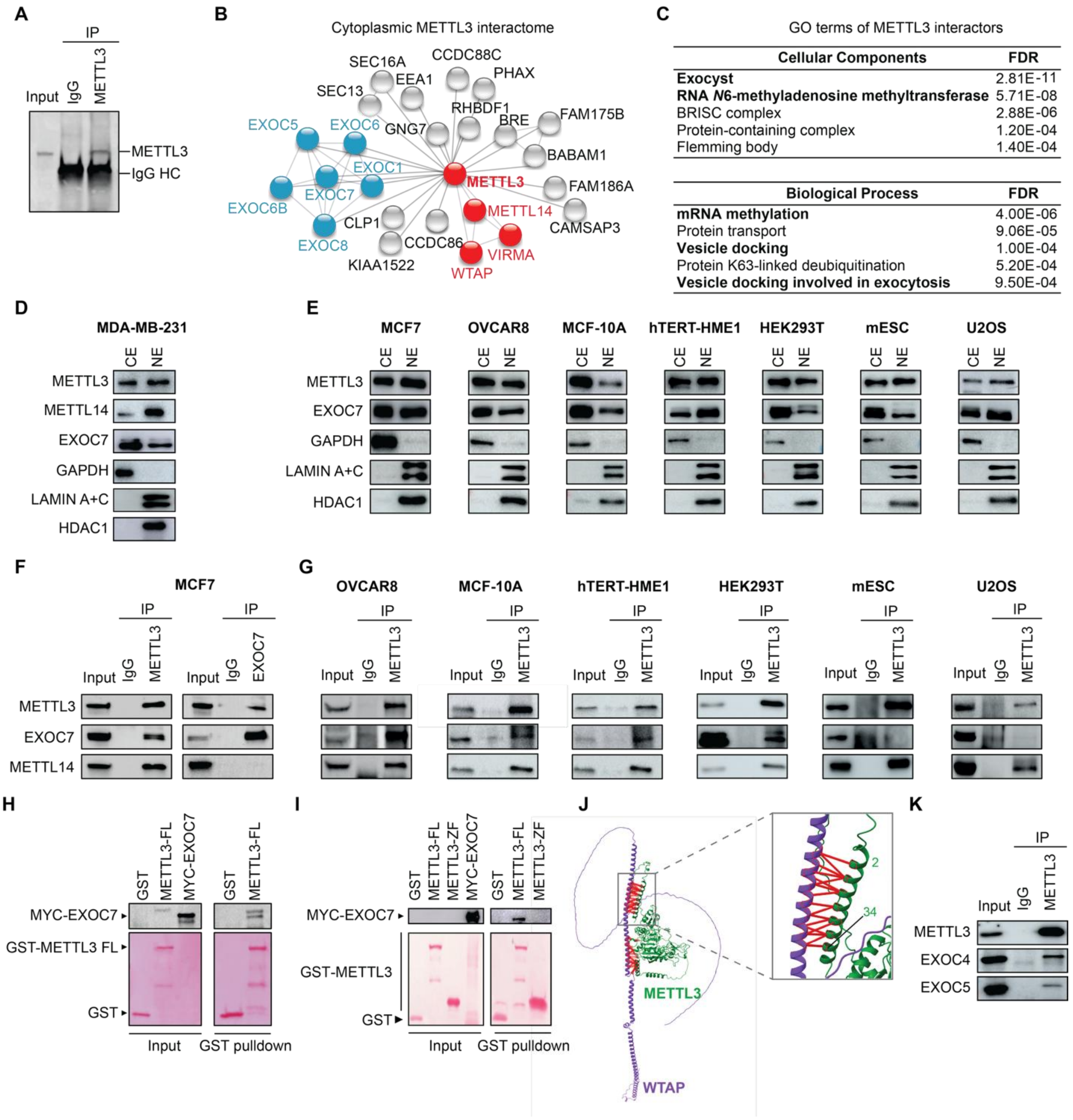


*Figure legend in the next page*

**Fig. S3. METTL3 cytoplasmic interactome and subcellular fractionation.** (**A**) Western blot of METTL3-IP from cytoplasmic extracts of MCF7 cells. IgG was used as control and 10% input was loaded. (**B**) STRING network of proteins identified by METTL3 immunoprecipitation from MCF7 cytoplasmic extracts followed by LC-MS/MS analysis. $m^6A$ regulators and exocyst subunits are highlighted. (**C**) Gene Ontology (GO) analysis of the cellular components (top) and biological processes (bottom) enriched among METTL3 cytoplasmic interactors identified by LC-MS/MS. (**D**) Western blot analysis of the subcellular localization of EXOC7, METTL3 and METTL14 in MDA-MB-231 cells. GAPDH, lamin A+C and HDAC1 were used as loading controls for the cytoplasmic (CE) and nuclear (NE) extracts. (**E**) Western blot analysis of METTL3 and EXOC7 from cytoplasmic and nuclear extracts of multiple cancer and non-transformed cell lines. GAPDH, lamin A+C and HDAC1 were used as loading controls. (**F** and **G**) Immunoprecipitation of cytoplasmic extracts using METTL3 and EXOC7 antibodies followed by immunoblotting with METTL3, EXOC7, and METTL14. IgG served as a control and 10% of input was loaded. (**H** and **I**) GST-pulldown assays of GST–METTL3 full-length (FL) and GST-METTL3-ZF with *in vitro* transcribed and translated MYC–EXOC7. The top panels show the blots of EXOC7 pulled down along with GST-tagged proteins. GST–tagged METTL3FL and METTL3-ZF were detected with Ponceau red staining. GST alone was used as the negative control. The percentage of input used is 1% for the MYC-EXOC7 and 10% for the GST-tagged proteins. (**J**) AlphaFold model illustrating the interaction of METTL3 and WTAP. Boundary residues of the interface are shown. (**K**) Immunoprecipitation of cytoplasmic extracts of MDA-MB-231 using METTL3 antibody followed by immunoblotting with METTL3, EXOC4 and EXOC5. IgG served as a control and 10% of input was loaded. Results are one representative of $n = 3$ (**D** to **E** and **H**) and $n = 2$ (**F**, **G** and **I**) independent biological experiments.

#### Figure S4

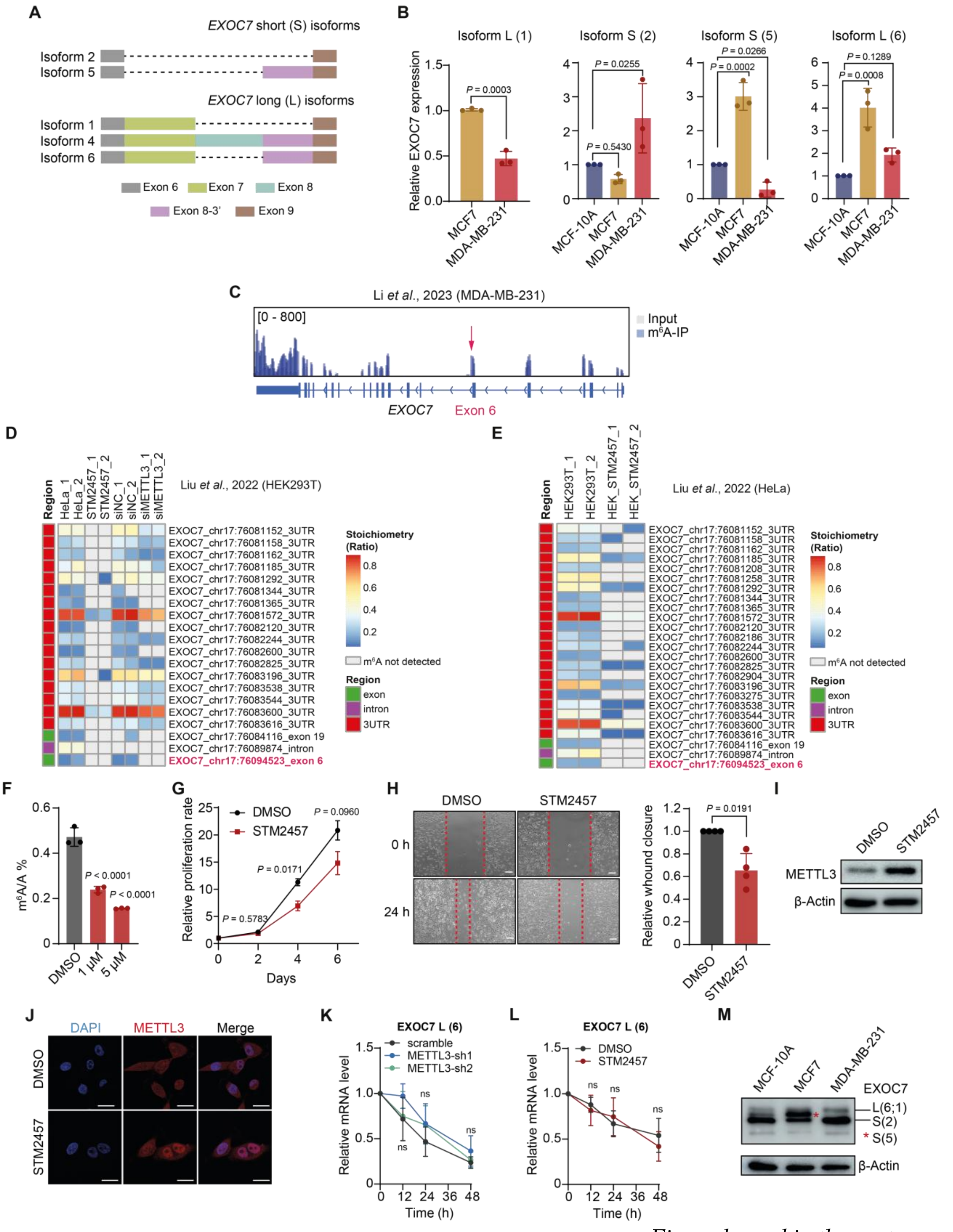


*Figure legend in the next page*

**Fig. S4. m$^6$A-dependent regulation of *EXOC7* Alternative Splicing (AS) and its functional impact in breast cancer cells.** (**A**) Schematic representation of *EXOC7* AS. The different exons are depicted in different colors. Isoforms are classified as short (S) or long (L) based on exclusion or inclusion of exon 7, respectively. (**B**) RT–qPCR analysis of *EXOC7* isoforms in MCF-10A, MCF7 and MDA-MB-231 cells. (**C**) m$^6$A peak distribution in *EXOC7* mRNA in MDA-MB-231 visualized in IGV. Input reads are represented in gray and enriched RNA immunoprecipitated in blue. Exon 6 is depicted in red. (**D** and **E**) Quantification of methylation levels for m$^6$A sites of *EXOC7* mRNA in HeLa and HEK293T using GLORI datasets. (**F**) LC-MS/MS quantification of m$^6$A in mRNA from MDA-MB-231 treated with DMSO (control) or with 1 and 5 μM of STM2457. (**G**) Relative cell proliferation rate of DMSO- and STM2457-treated MDA-MB-231 cells assessed over 6 days. (**H**) Left, wound healing assay. Right, quantification of gap closure quantification in DMSO- and STM2457-treated MDA-MB-231 cells after 24 h of treatment. Scale bars, 50 μm. (**I**) Western blot of METTL3 in MDA-MB-231 cells treated with DMSO or 10 μM STM2457. β-Actin is used as the loading control. (**J**) Immunofluorescence images of METTL3 in MDA-MB-231 treated with DMSO and 10 μM STM2457. DAPI was used as nuclear marker, respectively. Scale bars, 20 μm. (**K** and **L**) Line graphs depicting the effects of Actinomycin D treatment upon METTL3 depletion (**K**) and 10 μM STM2457 (**L**), on EXOC7 isoform 6 observed at 12, 24 and 48 h after treatment. (**M**) Western blot of EXOC7 isoforms in MCF-10A, MCF7, and MDA-MB-231 cells. The red asterisk indicates the intermediate band, corresponding to the short isoform 5. β-Actin is used as the loading control. Statistical analysis: Two-tailed Student's *t*-test (**B** (isoform 1), **G**, **H**, and **L**) and one-Way ANOVA with Dunnett's correction for multiple comparison (**B** (isoforms 2, 5 and 6), **F** and **K**). Data are mean ± SD; $n \geq 3$ (**B**, **F**, **G**, **H**, **K** and **L**). Results are one representative of $n = 4$ (**H** to **J**) and $n = 2$ (**M**) independent biological experiments.

**Figure S5**

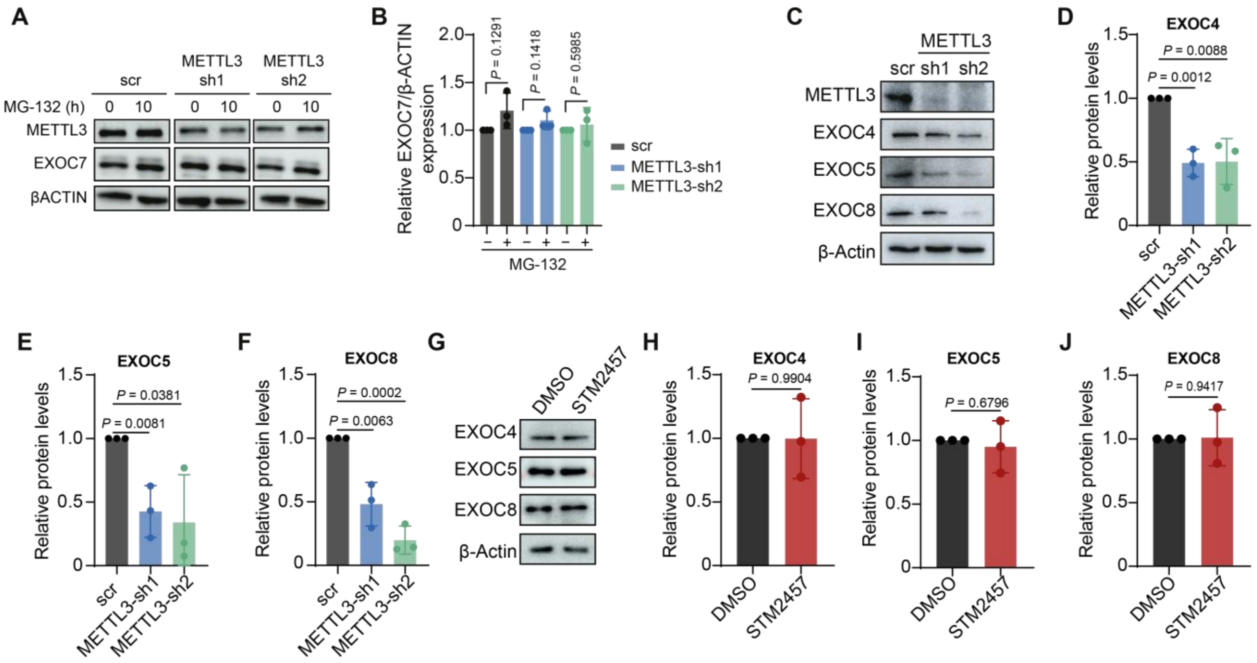


**Fig. S5. Proteasome inhibition does not affect EXOC7 levels.** (**A**) Western blot of METTL3 and EXOC7 at 0 and 10 h after MG-132 treatment. β-Actin is used as loading control. (**B**) Quantification of protein recovery of EXOC7 after quantification and normalization of bands from (**A**). (**C**) Western blot of METTL3, EXOC4, EXOC5, EXOC8 after METTL3 depletion. (**D** to **F**) Bar plots illustrating the protein levels of the exocyst components in (**C**). (**G**) Western blot of EXOC4, EXOC5, EXOC8 after DMSO or 10 μM STM2457. (**H** to **J**) Bar plots illustrating the protein levels of the exocyst components in (**G**). Statistical analysis: two-tailed Student's *t*-test (**B**, **D** to **F** and **H** and **I**). Data are mean ± s.d.; $n = 3$ (**B**, **D** to **F** and **H** and **I**). Results are one representative of $n = 3$ independent biological experiments (**A**, **C** and **G**).

#### Figure S6

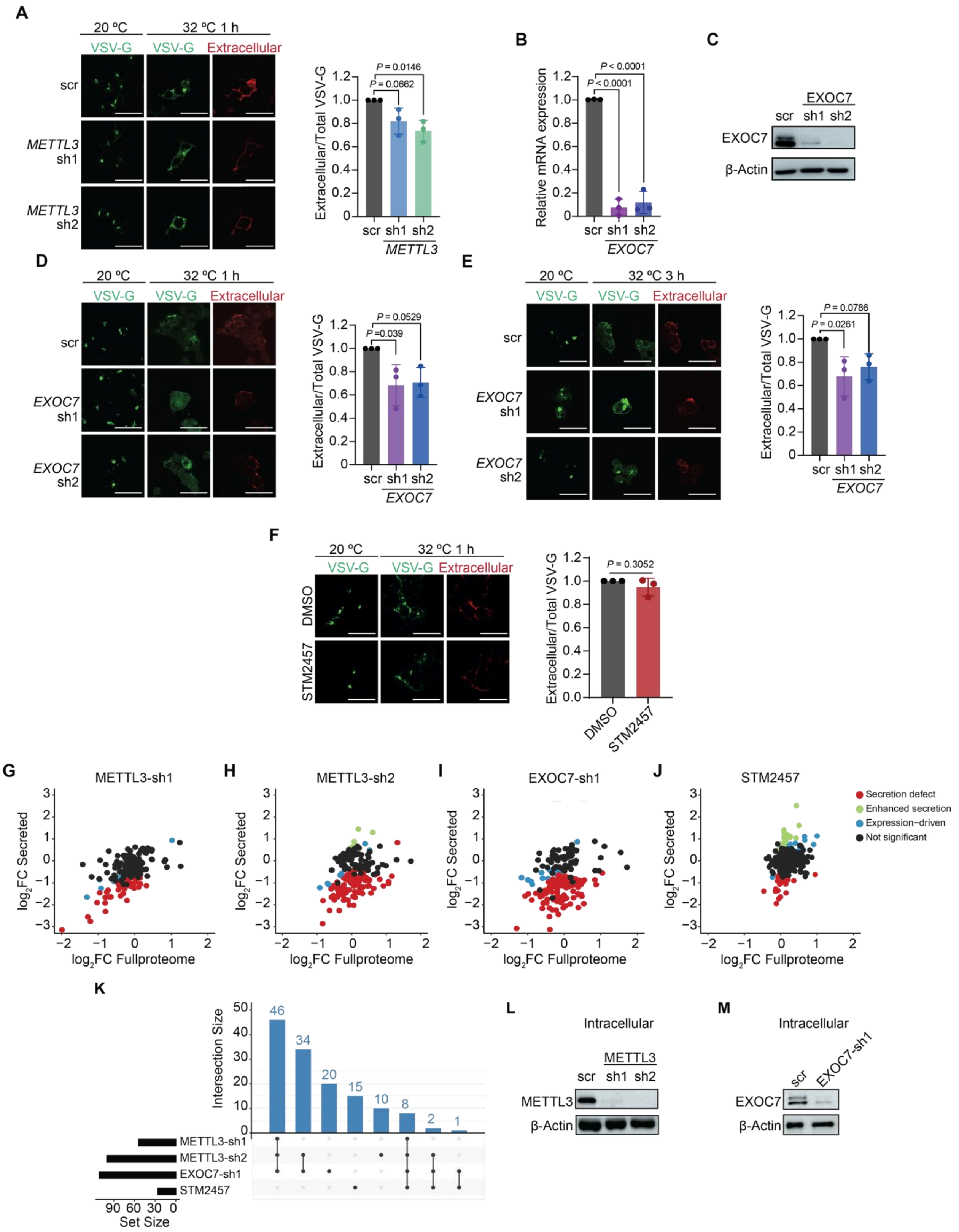


*Figure legend in the next page*

**Fig. S6. METTL3 modulates vesicle trafficking and the secretome.** (**A**) Left, immunofluorescence images of control (scramble (scr)) and *METTL3* knockdown (sh1 and sh2) HEK293T cells expressing GFP-VSV-Gts045 collected after 1 h at 32 ºC, then stained with the 8G5 antibody to visualize VSV-G incorporation into the plasma membrane. Right, quantification shows the ratio of extracellular (8G5 staining) to total cellular VSV-G fluorescence. (**B**) *EXOC7* mRNA levels and (**C**) protein expression upon depletion of *EXOC7* (sh1 and sh2) in MDA-MB-231 cells. β-Actin is used as the loading control. (**D**) Left, immunofluorescence images of scr and *EXOC7* knockdown (sh1 and sh2) HEK293T cells expressing GFP-VSV-Gts045 collected after 1 h at 32 ºC, immunostained with 8G5 antibody. Right, quantification shows the ratio of extracellular (8G5 staining) to total cellular VSV-G fluorescence. (**E**) Left, immunofluorescence images of scr and EXOC7 knockdown (sh1 and sh2) HEK293T cells expressing GFP-VSV-Gts045 collected after 3 h at 32 ºC, immunostained with 8G5 antibody. Right, quantification shows the ratio of extracellular (8G5 staining) to total cellular VSV-G fluorescence. (**F**) Left, immunofluorescence images of HEK293T cells expressing GFP-VSV-Gts045 and treated with DMSO (control) or STM2457 inhibitor after 1 h at 32 ºC. Right, quantification shows the ratio of extracellular (8G5 staining) to total cellular VSV-G fluorescence. (**G** to **J**) Scatter plots comparing protein abundance in the full proteome (x-axis) and secretome (y-axis) for METTL3-sh1, METTL3-sh2, EXOC7-sh1, and STM2457 versus respective controls. Proteins with secretion defect (red), enhanced secretion (green), and expression-driven secretion (blue) are highlighted. Non-significant proteins are shown in black. (**K**) UpSet plot showing the number of common and unique downsecreted proteins in METTL3 knockdowns, EXOC7-sh1 and cells treated with STM2457. (**L** and **M**) Western blot showing knockdown efficiency of METTL3 (**L**) and EXOC7 (**M**) in intracellular protein extracts prepared for secretome analysis. Statistical analysis: One-Way ANOVA with Dunnett's correction for multiple comparison (**A**, **B**, **D** and **E**), and two-tailed Student's t-test (**F**). Data are mean ± SD; $n = 3$ (**A**, **B**, **D** to **F**). Results are one representative of $n = 3$ independent biological experiments (**C**, **L** and **M**). Scale bars, 30 µm.

#### Figure S7

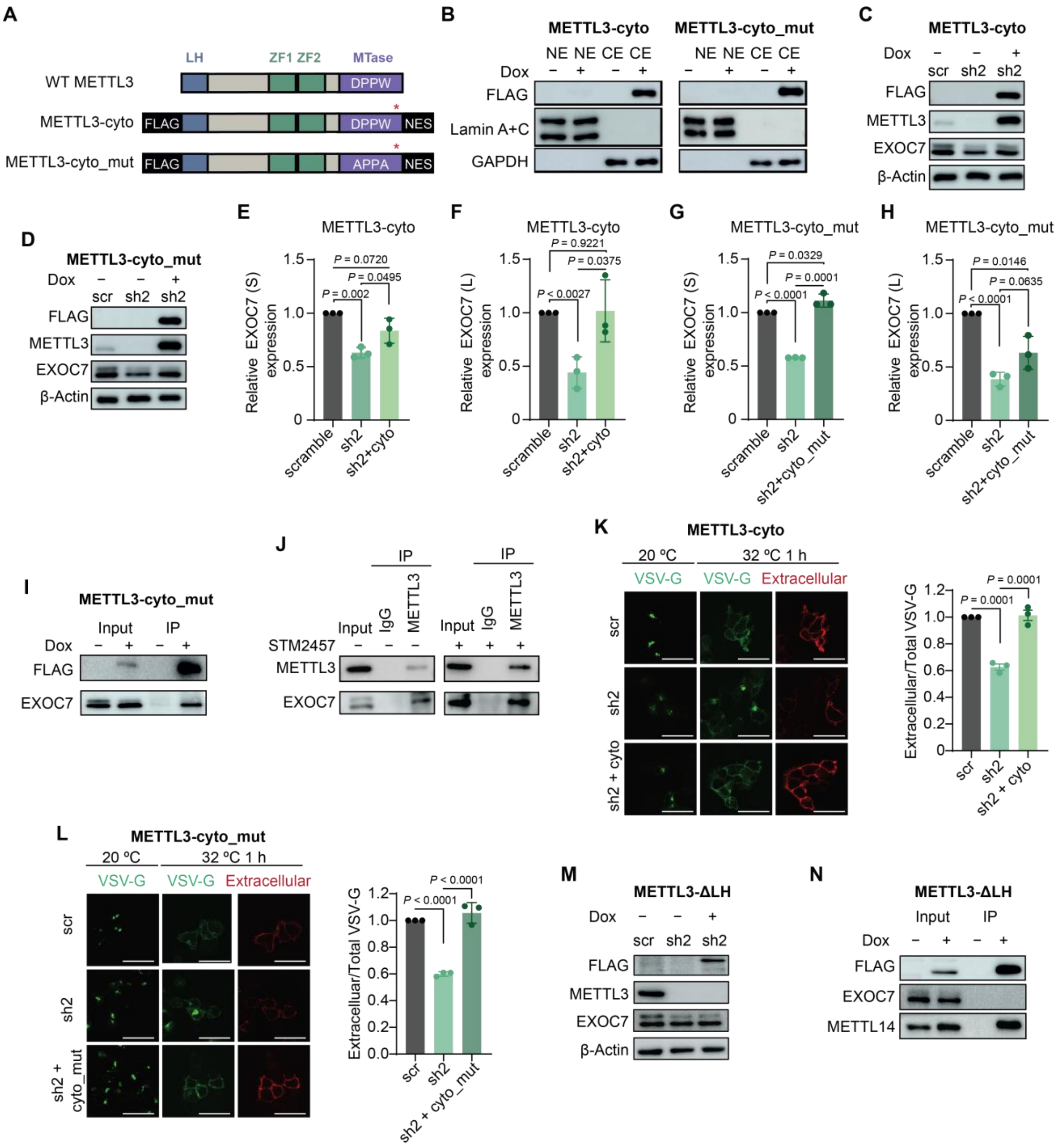


*Figure legend in the next page*

**Fig. S7. METTL3 rescue strategy and validation assays.** (**A**) Schematic of wild-type METTL3 (WT), doxycycline-inducible FLAG-tagged cytoplasmic METTL3 (METTL3-cyto), and catalytically inactive METTL3-cyto_mut (DPPW→APPA: D395A+W398A). A nuclear export signal (NES) was added to maintain cytoplasmic localization. METTL3 domains are indicated: leader helix (LH), zinc fingers (ZF1, ZF2), and methyltransferase (MTase). Red asterisks mark silent mutations at shRNA2 sites enabling inducible expression. (**B**) Western blot showing subcellular localization of FLAG-tagged METTL3 variants; GAPDH and lamin A+C serve as cytoplasmic (CE) and nuclear (NE) controls. (**C** and **D**) Western blots of FLAG, METTL3, and EXOC7 in scramble (scr) and METTL3-knockdown (sh2) cells, and after expression of WT METTL3-cyto (**C**) or METTL3-cyto_mut (**D**). β-Actin is the loading control. (**E** and **F**) Quantification of protein EXOC7 levels for the short (**E**) and long (**F**) isoforms after quantification and normalization of bands from (**C**). (**G** and **H**) Quantification of protein EXOC7 levels for the short (**G**) and long (**H**) isoforms after quantification and normalization of bands from (D). (**I**) FLAG immunoprecipitation from METTL3-cyto_mut cytoplasmic extracts showing METTL3–EXOC7 interaction (10% input). (**J**) METTL3 immunoprecipitation from DMSO- and STM2457-treated cells showing METTL3–EXOC7 interaction. (**K** and **L**) Immunofluorescence of HEK293T cells expressing GFP-VSV-Gts045 after 1 h at 32 °C, stained with 8G5 antibody to visualize plasma membrane incorporation. (**K**) Rescue with METTL3-cyto; (**L**) rescue with METTL3-cyto_mut. Right: quantification of extracellular/total VSV-G fluorescence ratio. (**M)** Western blot of FLAG, METTL3, and EXOC7 in scr, sh2, and METTL3-ΔLH rescue cells. METTL3 antibody does not detect METTL3-ΔLH (epitope within first 34 residues). β-Actin is the loading control. (**N**) FLAG immunoprecipitation from METTL3-ΔLH cytoplasmic extracts showing no EXOC7 interaction. Input loaded is 10%. Statistical analysis: Two-tailed Student's *t*-test (**E** to **H**, **K** and **L**). Data are mean ± SD; $n = 3$ (**E** and **H**, **K** and **L**). Results are one representative of $n = 2$ (**B** and **I**) and $n = 3$ (**C**, **D**, **J**, **M** and **N**) independent biological experiments (**C**, **L** and **M**). Scale bars, 30 μm.

### Figure S8

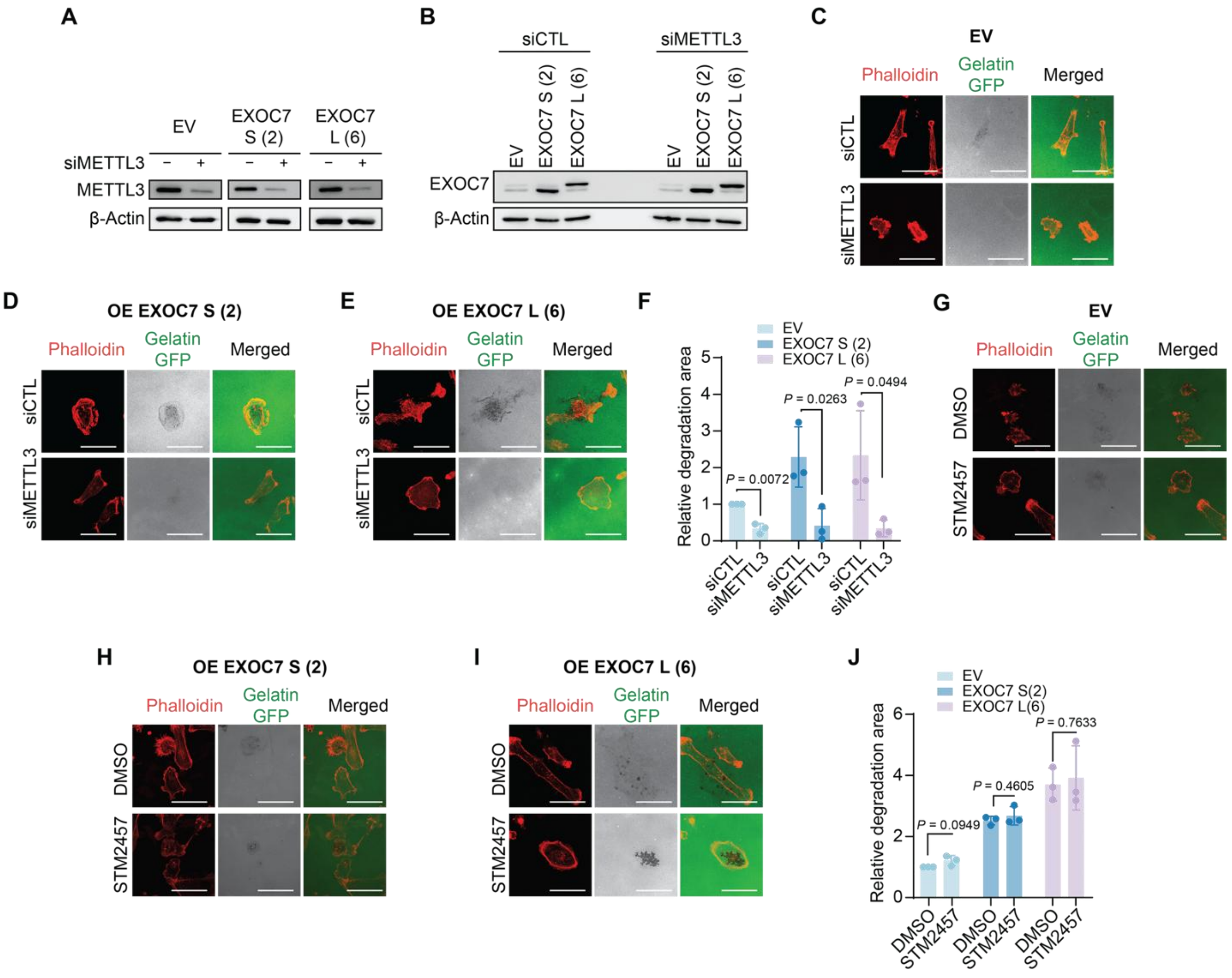


**Fig. S8. Effect of EXOC7 overexpression in invadopodia formation of MDA-MB-231 cells.** (**A**) Western blot showing METTL3 levels in cells transfected with empty vector (EV), EXOC7 short isoform 2 (S) and EXOC7 long isoform 6 (L), treated with siRNA control (siCTL) and siRNA against METTL3 (siMETTL3). β-Actin serves as loading control. (**B**) Western blot showing EXOC7 expression in cells transfected with empty vector (EV), EXOC7 S and EXOC7 L with or without METT3 silencing. β-Actin serves as loading control. (**C** to **E**) Gelatin degradation assay of MDA-MB-231 treated with siCTL and siMETTL3 upon overexpression of empty vector (EV) (**C**), EXOC7 S (**D**) and EXOC7 L (**E**). (**F**) Quantification of areas of degradation corresponding to invadopodia formation in (**C** to **E**). (**G** to **I**) Gelatin degradation assay of MDA-MB-231 treated with DMSO or STM2457 and upon overexpression of empty vector (EV) (**G**), EXOC7 S (**H**) and EXOC7 L (**I**). (**J**) Quantification of areas of degradation corresponding to invadopodia formation in (**G-I**). Statistical analysis: Two-tailed Student's *t*-test (**F** and **J**). Data are mean ± SD; $n = 3$ (**F** and **J**). Results are one representative of $n = 3$ (**A** and **B**) independent biological replicates. Scale bars, 40 μm.

#### Figure S9

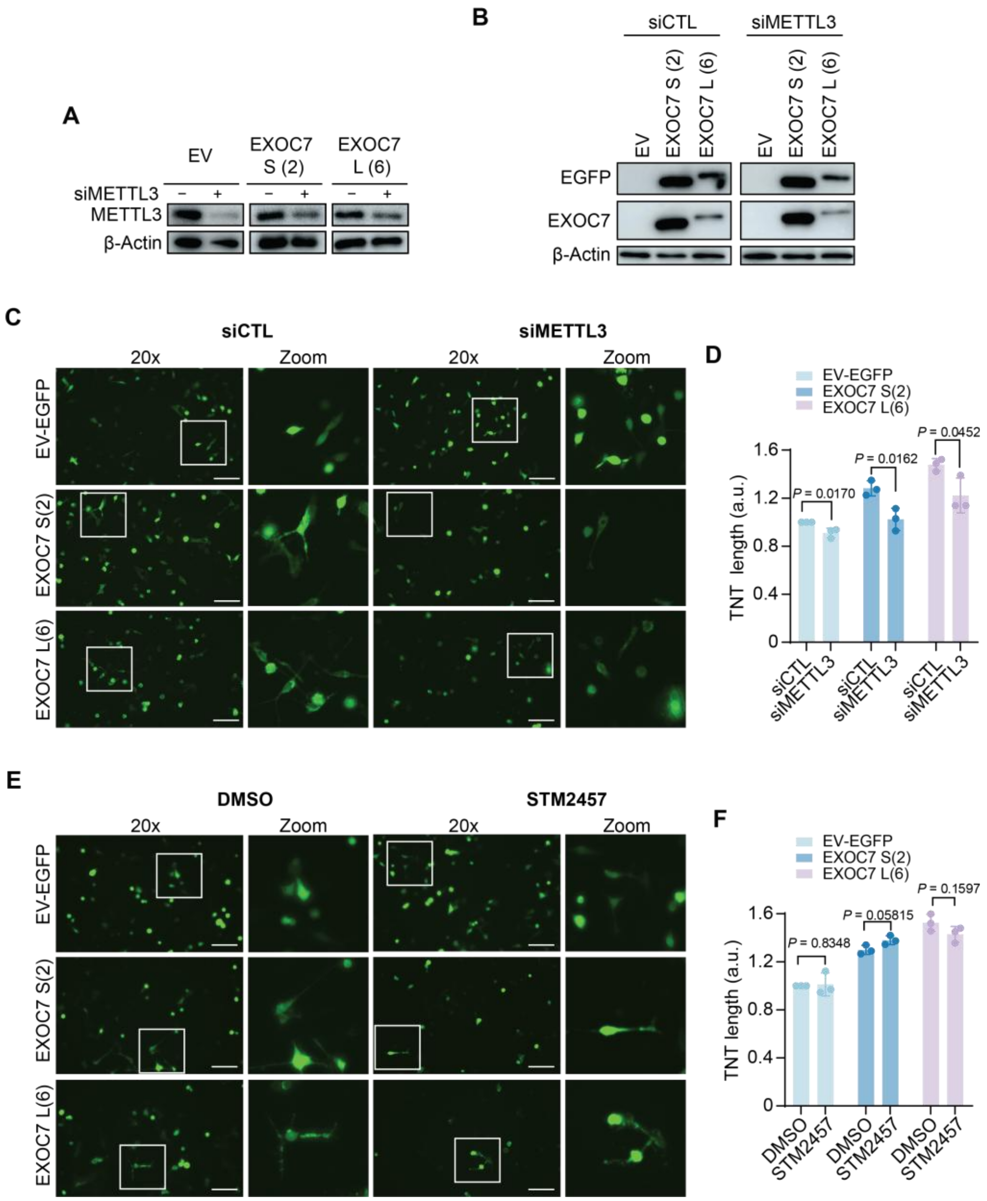


*Figure legend in the next page*

**Fig. S9. Effect of EXOC7 overexpression in TNTs of MDA-MB-231 cells.** (**A**) Western blot showing METTL3 levels in cells transfected with empty pEGFP plasmid (EV-EGFP), pEGFP-C3-EXOC7 short isoform 2 (S) or long isoform 6 (L), under siCTL and siMETTL3 conditions. β-Actin serves as loading control. (**B**) Western blot of EXOC7 and EGFP in cells transfected with empty pEGFP plasmid (EV-EGFP), pEGFP-C3-EXOC7 S or L isoforms, showing proper EXOC7 isoform overexpression, under siCTL and siMETTL3 conditions. β-Actin is used as the loading control. **(C)** Fluorescence images of cells treated with siCTL or siMETTL3 transfected with empty pEGFP plasmid (EV–EGFP), pEGFP–C3–EXOC7 S or L isoforms. **(D)** Quantification of TNT length from (**C**); a.u., arbitrary units. **(E)** Fluorescence images of cells treated with DMSO (control) or STM2457 transfected with empty pEGFP plasmid (EV–EGFP), pEGFP–C3–EXOC7 S or L isoforms. **(F)** Quantification of TNT length from (**E**). a.u., arbitrary units. Statistical analysis: Two-tailed Student's *t*-test (**D** and **F**). Data are mean ± SD; $n = 3$ (**D** and **F**). Results are one representative of $n = 3$ (**A** and **B**) independent biological replicates. Scale bar, 200 μm.

**Table S4. Primers, shRNAs and gRNAs used in this study.**

| Name | Application | Sequence |
|---|---|---|
| Actin_Forward | RT-qPCR | AGATCAAGGTGGGTGTCTTTC |
| Actin_Reverse | RT-qPCR | AGCAATGATCTGAGGAGGGAAG |
| METTL3_Forward | RT-qPCR | AACTGCAACGCATCATTCGG |
| METTL3_Reverse | RT-qPCR | TTGACACCAACCAAGCAGTG |
| EXOC7-iso1-Forward | RT-qPCR | TTCCTCTGGAAGGGAGAGATGAC |
| EXOC7-iso1-Reverse | RT-qPCR | GGCATTGCTGGTGAGCTCGTGTA |
| EXOC7-iso2-Forward | RT-qPCR | CAAGCGGCCAGGGAGAGATGAC |
| EXOC7-iso2-Reverse | RT-qPCR | GGCATTGCTGGTGAGCTCGTGTA |
| EXOC7-iso5-Forward | RT-qPCR | AGCGGCCAGGTCACGAGC |
| EXOC7-iso5-Reverse | RT-qPCR | TCGGACAGGTGCTTAACTCGGAAAT |
| EXOC7-iso6-Forward | RT-qPCR | TGGCCGCAACCAAGATTTCATG |
| EXOC7-iso6-Reverse | RT-qPCR | ATGCTCGTGACCTTCCAGAG |
| EXOC7 iso5+iso6 Forward | RT-PCR | TGGCCGCAACCAAGATTTCATG |
| EXOC7 iso5+iso6 Forward | RT-PCR | TCGGACAGGTGCTTAACTCGGAAAT |
| EXOC7 all isoforms-Forward | RT-qPCR | TGGCCGCAACCAAGATTCATG |
| EXOC7 all isoforms-Reverse | RT-qPCR | GAGAAGTCGTGTCGCACAATGGC |
| pLKO.1-shMETTL3_1 | sh1 | GCAAGTATGTTCACTATGAAA |
| pLKO.1-shMETTL3_2 | sh2 | CGTCAGTATCTTGGGCAAGTT |
| pLKO.1-shMETTL3_3 | sh3 | GCTAAACCTGAAGAGTGATAT |
| pLKO.1-shEXOC7_1 | sh1 | CCTGCACAACAACTACAATTA |

| | | |
|---|---|---|
| pLKO.1-shEXOC7_2 | sh2 | GCTGCAGGAGAATGTTGAGAA |
| GGG+XhoI+EXOC7 Forward | Cloning EXOC7 isoforms in pEGFP-C3 | GGGCTCGAGATGATTCCCCCACAGG AGGCATCCGC |
| GGG+BamHI+EXOC7 Reverse | Cloning EXOC7 isoforms in pEGFP-C3 | GGGGGATCCTCAGGCAGAGGTGTCG AAAAGGCGA |
| GGG+_NdeI +MYC+EXOC7 forward | Cloning Myc-EXOC7 in pET22b+ | GGGCATATGGAACAGAAACTGATTA GCGAAGAAGACCTGATGATTCCCCC ACAGGAGGC |
| GGG+EcoRI+EXOC7 reverse | Cloning Myc-EXOC7 in pET22b+ | GAATTCTCAGGCAGAGGTGTCGAAA A |
| METTL3-Forward | Cloning full length METTL3 in pGEX6P-2 | TTCCAGGGGCCCCTGGGATCAATGT CGGACACGTGGAGCTCT |
| METTL3-Reverse | Cloning full length METTL3 in pGEX6P-2 | GATGCGGCCGCTCGACTCGACTTAT AAATTCTTAGGTTTAGAGATGATA |
| METTL3_LH_Forward | Cloning LH domain METTL3 in pGEX6P-2 | TTCCAGGGGCCCCTGGGATCAATGT CGGACACGTGGAGCTCT |
| METTL3_LH_Reverse | Cloning LH domain METTL3 in pGEX6P-2 | GATGCGGCCGCTCGACTCGAC TTAATCCAAGTGCCCCGAGT |
| METTL3_ZF_Forward | Cloning zinc finger domain METTL3 in pGEX6P-2 | TTCCAGGGGCCCCTGGGATCA<br><br>TCCATTGTTGAAAAATTTCG |
| METTL3_ZF_Reverse | Cloning zinc finger domain METTL3 in pGEX6P-2 | GATGCGGCCGCTCGACTCGAC TTAGCAAGCATCAATTTCAT |
| METTL3_NT_Forward | Cloning N-terminal domain METTL3 in pGEX6P-2 | TTCCAGGGGCCCCTGGGATCAATGT CGGACACGTGGAGCTCT |
| METTL3_NT_Reverse | Cloning N-terminal domain METTL3 in pGEX6P-2 | GATGCGGCCGCTCGACTCGAC TTAGAGTCGGTCTGCACTGG |
| METTL3_MT_Forward | Cloning Methyl transferase domain METTL3 in pGEX6P-2 | TTCCAGGGGCCCCTGGGATCATTCC CACCTCAGTGGATCTG |

| | | |
|---|---|---|
| METTL3_MT_Reverse | Cloning Methyl transferase domain METTL3 in pGEX6P-2 | GATGCGGCCGCTCGACTCGACTTAT AAATTCTTAGGTTTAG |
| METTL3-cyto Forward | Cloning METTL3 in pInd20 | CCGTCAGATCGCCTGACCATGGATT ACAAGGATGACGATGACAAGTCGG ACACGTGGAGCTCTAT |
| METTL3-cyto_NES_Reverse | Cloning METTL3 in pInd20 | GCCCTCTAGACTCGACTAGTCCAGG GTCAGGCGCTCCAGGGGAGGCAGTT GCAGTAAATTCTTAGGTTTAGAGA |
| METTL3-ΔLH Forward | Cloning LH domain METTL3 in pInd20 | CCGTCAGATCGCCTGACCATGGATT ACAAGGATGACGATGACAAGTCGG ACACGTGGAGCTCTAT |
| METTL3-ΔLH Reverse | Cloning LH domain METTL3 in pInd20 | GCCCTCTAGACTCGACTAATCCAAG TGCCCCGAGT |
| TCC+BamHI+EXOC7 Forward | Cloning EXOC7 isoforms in pCDNA3 | TCCGGATCCATGATTCCCCCACAGG AGGCA |
| TCC+EcoRI+EXOC7 Reverse | Cloning EXOC7 isoforms in pCDNA3 | TCCGAATTCtcaggcagaggtgtcgaaaag |

**Table S5. Criteria used to define categories in the secretome analysis.**

| Category | Δ Secretion Efficiency (log2FC secreted − log2FC fullproteome) | FDR (Secreted) | Additional Criteria | Biological Interpretation |
|---|---|---|---|---|
| Secretion defect | ≤ -0.5 | < 0.05 | — | Significant reduction in secretion not explained by expression |
| Enhanced secretion | ≥ 0.5 | < 0.05 | log2FC_secreted > 0 | Significant increase in secretion |
| Expression-driven / No secretion-specific change | > -0.5 and < 0.5 OR ≤ -0.5 but log2FC_secreted < 0 & log2FC_fullproteome < 0 | < 0.05 | — | Changes explained by overall protein expression, not secretion |
| Not significant | Any | ≥ 0.05 | — | No statistically significant change |

**Tables included as Excel spreadsheets:**

**Table S1. Mass spectrometry of METTL3-interacting proteins.** Number of unique peptides, sequence coverage, p-value, name, and accession number are indicated.

**Table S2. Mass spectrometry of secretome analysis for METTL3 and EXOC7 knockdowns.** Experimental condition, Uniprot ID, log2FC_secreted, FDR_secreted, log2C_fullproteome, FDR_fullproteome, delta_secretion_efficiency and category are indicated.

**Table S3. Mass spectrometry of secretome analysis for STM2457 treatment.** Experimental condition, Uniprot ID, log2FC_secreted, FDR_secreted, log2C_fullproteome, FDR_fullproteome, delta_secretion_efficiency and category are indicated
